# GNN4DM: A Graph Neural Network-based method to identify overlapping functional disease modules

**DOI:** 10.1101/2024.01.30.578064

**Authors:** András Gézsi, Péter Antal

**Affiliations:** Department of Measurement and Information Systems, Budapest University of Technology and Economics, Magyar tudosok krt. 2. Budapest, H-1117, Hungary

**Keywords:** Disease module identification, Graph Neural Network, Overlapping community detection

## Abstract

**Motivation:** Identifying disease modules within molecular interaction networks is an essential exploratory step in computational biology, offering insights into disease mechanisms and potential therapeutic targets. Traditional methods often struggle with the inherent complexity and overlapping nature of biological networks, and they are limited in effectively leveraging the vast amount of available genomic data and biological knowledge. This limitation underscores the need for more effective, automated approaches to integrate these rich data sources.

**Results:** In this work, we propose GNN4DM, a novel graph neural network-based structured model that automates the discovery of overlapping functional disease modules. GNN4DM effectively integrates network topology with genomic data to learn the representations of the genes corresponding to functional modules and align these with known biological pathways for enhanced interpretability. Following the DREAM benchmark evaluation setting and extending with three independent data sources (GWAS Atlas, FinnGen, and DisGeNET), we show that GNN4DM performs better than several state-of-the-art methods in detecting biologically meaningful modules. Furthermore, we highlight the method’s applicability through its discovery of two novel multimorbidity-modules significantly enriched in a wide array of apparently unrelated diseases.

**Availability and Implementation:** Source code, all training data, and all identified disease modules are freely available for download at https://github.com/gezsi/gnn4dm. GNN4DM was implemented in Python.

## Introduction

It’s widely recognized that molecular networks exhibit a high degree of modularity, meaning, there exist subgroups of nodes that are more densely interconnected than would be anticipated by random chance. Often, these individual modules encapsulate genes or proteins that partake in identical or closely related biological functions (1, 2). This characteristic has been widely leveraged to infer unknown gene functions, a principle termed “guilt-by-association” (3). Similarly, genes associated with the same disease or symptom tend to cluster together in molecular networks, as demonstrated in various studies (4–6).

By convention, a *disease module* is defined as a topological network module associated with a disease, specifically a network module whose malfunctioning components (genes or proteins) contribute to the abnormal phenotype associated with that disease (7). Identifying disease modules within molecular interaction networks has contributed to uncovering the genetic basis of a wide range of complex diseases, offering insights into their underlying molecular mechanisms (8, 9), facilitating the identification of potential therapeutic targets (10), and in various other tasks, such as quantifying the associations between diseases (11).

Traditionally, disease module identification aims to uncover cohesive modules (also known as communities or clusters) with denser internal connections compared to external ones. However, identifying disease modules is an inherently ill-posed problem, as the interactome, similarly to most real networks, lacks a clear community structure, yet it is characterized by well-defined statistics of overlapping and nested communities (12).

In this study, we adopt a more integrative approach towards modules, targeting the identification of *functional disease modules* that are not merely cohesive but also collectively contribute to higher-level biological functions reflecting the modular and hierarchical organization observed in biological systems. Our proposed method hinges on two core modeling assumptions, namely:

*Assumption 1: Functional disease modules are overlapping*. Functional modules in biological networks often exhibit a significant degree of overlap. This overlapping nature is indicative of the multi-functionality of genes, where a single gene can be part of multiple functional groups or pathways. This assumption is well-acknowledged, with many studies reporting an overlapping community structure across various biological networks (7, 13).

*Assumption 2: Functional modules are organized into a complex, hierarchical system that encompasses known biological pathways*. We posit that either 1) biological pathways are orchestrated by smaller, cohesive functional modules, or 2) these pathways themselves interact, potentially serving as components of even larger functional modules. In either case, the functional modules represent clusters of genes working together to execute specific biological functions. This modular organization is not only a hallmark of biological complexity but also a manifestation of functional redundancy and robustness in biological systems (14–18).

In this work, we propose a deep learning-based method, termed Graph Neural Network for Disease Modules (GNN4DM), that leverages these assumptions to identify overlapping functional disease modules that align well with known biological pathways. By employing a two-step approach, initially, a Graph Convolutional Network (GCN) is utilized to fuse network data and genomic data (comprising gene expression and genome-wide association data) to generate a community affiliation matrix representing the likelihood of each gene belonging to various modules. Subsequently, the identified modules are refined by leveraging known pathway information from various pathway databases utilizing constrained logistic regression models, aiming to align the modular structure with established biological knowledge. The two components of the model are trained end-to-end to derive interpretable functional disease modules.

We assessed the model’s ability to identify biologically relevant modules using three large-scale independent data sources (GWAS Atlas, FinnGen and DisGeNET), examining their association with complex traits and diseases. Our findings reveal that GNN4DM outperforms several widely used and state-of-the-art methods in detecting a high proportion of biologically meaningful modules associated with a broad spectrum of complex traits. Additionally, we highlight the interpretability of the identified modules by presenting two highly enriched modules.

### Background work

Disease module identification approaches are broadly categorized into general and disease-specific methods. General methods aim to reveal disease modules based on the topology of molecular networks without tailoring them to any particular disease. This involves identifying locally dense *communities* within a network. In contrast, disease-specific methods aim to uncover disease-pertinent modules employing disease-specific data, like genome-wide association signals or cancer mutation data. Our proposed approach aligns with general methods (see Manipur et al. (19) for a recent review).

This study primarily focuses on methods based on graph neural networks (GNNs). Recently, GNNs have emerged as effective tools for leveraging graph-structured data. They efficiently capture the relationships among nodes within a network while also incorporating node attributes, achieving state-of-the-art results in various tasks.

NOCD, introduced by Shchur and Günnemann (20), was one of the pioneering neural models for overlapping community detection. It combines the power of GNNs with the Bernoulli–Poisson graph generative probabilistic model. This model learns a non-negative community affiliation matrix through a GNN, which represents the nodes’ assignment into communities. Our proposed method builds on this model, extending it by fine-tuning the module representations to better align with existing biological knowledge.

Deepgmd uses a similar formalism to NOCD aiming to detect gene regulator modules within gene co-expression networks derived from gene expression profile data (21). While it uses a similar graph generative model for community affiliations, Deepgmp replaces the Poisson model with a sigmoid function that is used to translate the probability of two nodes sharing a community into the probability of an edge existing between them.

UCoDe, introduced by Moradan et al. (22), performs both overlapping and non-overlapping community detection. It utilizes a contrastive loss function that maximizes a soft version of network modularity, offering a unified approach to community detection in networks.

## Materials and methods

### Datasets

The architectural design of our model is based on using two different types of datasets. Primarily, datasets encompassing genome-wide information, along with structural properties of the nodes in the underlying graph, were utilized as input features in the model. Conversely, annotations from pathway databases possessing partial, specialized, and curated information were leveraged as auxiliary tasks for the model to predict.

### Data source for graph structure

#### Protein-protein interactions

We utilized protein-protein interaction data from STRING-DB v12.0 (23), following a comprehensive study by Huang et al. (24) that assessed 21 human tissue-unaware interaction networks for their capability in predicting disease genes, and identified STRING as one of the top-performing networks. We mapped the protein identifiers to their corresponding Ensembl gene identifiers using the Ensembl Biomart tool. We selected only the highly confident interactions by filtering them based on their combined score, keeping those above 0.7. The resulting pairwise interactions defined the structure of the unweighted graph in which we identified the modules. The graph nodes represented 15793 protein-coding genes, and the edges denoted 234179 functional interactions between these genes.

### Data sources used for constructing input features

#### Gene expression measurement data

We used genomewide gene expression measurement data from the Genotype-Tissue Expression (GTEx) project (25). We used the bulk RNA-seq tissue expression dataset from the GTEx Analysis V8, which contains median gene-level transcript per million (TPM) values across 54 human tissues. We log-transformed the raw TPM values and imputed the missing expression values for 218 genes absent in the GTEx dataset by the tissue-wise mean TPM values. We used the transformed expression values as part of the input features for the gene nodes.

#### Gene-level genome-wide association data

We utilized gene-level genome-wide association data from the GWAS Atlas project Release 3 (26), which encompasses 4756 publicly available GWAS summary statistics. We downloaded the dataset containing the gene-level p-values computed by the MAGMA software for 19436 protein-coding genes. We split the dataset into two parts; 3545 summary statistics exclusive to studies other than UKBiobank were employed as inputs, while the 1211 UKBiobank-specific summary statistics were utilized for evaluation purposes (see Evaluation section). We imputed the missing p-values for 519 genes absent in the dataset with 0.5. We computed the negative log-transformed p-values (the higher, the more significant) and performed a PCA transformation for dimensionality- and noise reduction of the resulting significance values. We used the first 512 transformed dimensions and used these as part of the input features for the gene nodes. The total variance explained by the 512 principal components was 0.718.

#### Centrality measures of nodes within the network

Additionally, we calculated five other metrics using the `NetworkX` Python package (27) to quantify the centrality or importance of nodes within the network. These consist of degree, betweenness, eigenvector, and closeness centrality, and the PageRank score.

All the above vectors were concatenated and standardized to form 571-dimensional input feature vectors for the gene nodes.

#### Pathway databases

We derived gene sets from the MSigDB v2023.1 pathway datasets (28), namely the BioCarta, KEGG, Reactome, and WikiPathways databases encompassing canonical representations of biological processes compiled by domain experts. We filtered all datasets to those pathways containing at least 10 but no more than 500 genes in the graph. This resulted in 221 pathways for BioCarta (comprising 1324 unique genes), 186 pathways for KEGG (4889 genes), 1288 for Reactome (9905 genes), and 621 pathways for WikiPathways (7166 genes).

The pathways in these datasets served as auxiliary tasks that the model was trained to predict.

### The GNN4DM framework

We created an interpretable, graph neural network-based framework called GNN4DM that is used to identify overlapping functional disease modules in a protein-protein interaction network. The overview of the method is shown in Figure 1. The model is structured into two components. The initial component processes the underlying protein-protein interaction graph along with the genes’ input features, employing graph convolution layers to derive an inner representation for the nodes that correspond to their community affiliations. The subsequent component leverages the biological knowledge available in various pathway databases as auxiliary tasks, fine-tuning the inner representations of the genes such that the modules align with the structured biological knowledge in an interpretable way.

**Fig. 1.**
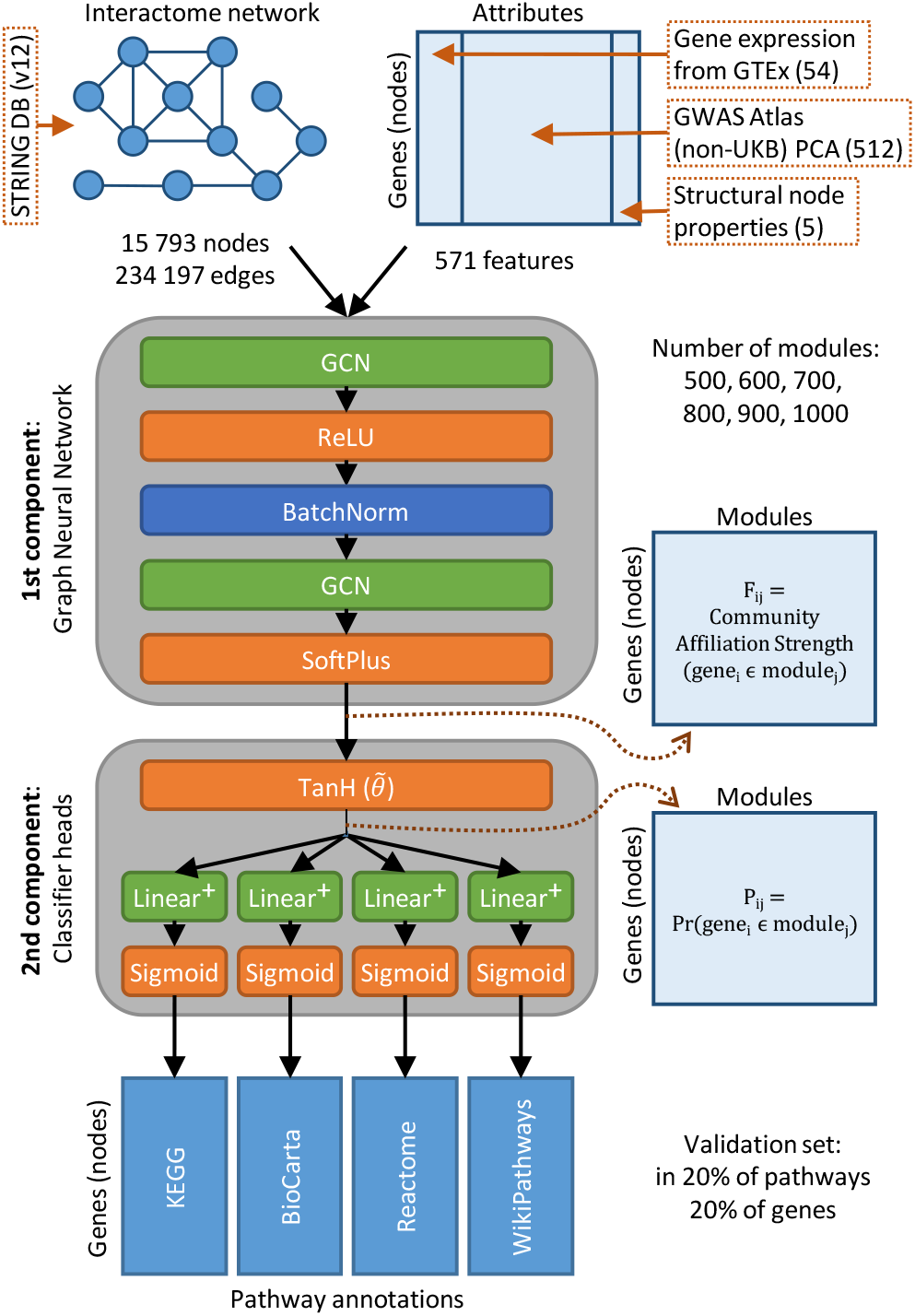
The GNN4DM framework. Datasets used for constructing the human interactome network (graph) and the attributes of the gene nodes in the graph are shown on top. The first component of the framework employs a graph neural network to process the underlying graph and the genes’ input features to derive an inner representation for the nodes corresponding to their non-negative community affiliation strengths. The second component of the framework refines the inner representations (converted first into module membership probabilities) of the genes such that the identified modules align with biological pathways contained in the KEGG, Bio-Carta, Reactome, and WikiPathways databases. This second component employs a constrained multivariable logistic regression to perform binary classification over each pathway in each pathway database. GCN: Graph Convolutional Network layer.

### Notations

Assuming an undirected, unweighted, static graph *G*(*V, E*) consisting of nodes *V* corresponding to human protein-coding genes and edges *E* representing the functional interactions between the corresponding gene nodes, we denote the binary adjacency matrix of *G* as **A** ∈{0, 1} ^*N* ×*N*^, where N is the number of nodes *V* = {1, …, *N* }, and *E* ={ (*u, v*) ∈ ∈*V*×*V* : *A*_*uv*_ = 1} is the set of edges in the graph. Every node is associated with a D-dimensional input vector that is represented as a feature matrix **X** ∈ ℝ^*N* ×*D*^ for all nodes.

### Identifying the overlapping modules via Graph Convolutional layers

The main aim of the GNN4DM framework is to find overlapping modules in the graph. This is achieved by assigning the gene nodes into *M* number of modules where the assignment is represented by a non-negative *community affiliation matrix* 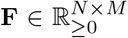, where *F*_*um*_ represents the belongs to module *strength* by which node *u* belongs to module *m*.

The first component of the GNN4DM framework uses Graph Convolutional Network layers (29) to generate the **F** community affiliation matrix, namely:

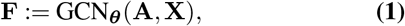

where ***θ*** represents the parameters of these layers. In our model, we used two GCN layers, applying a ReLU activation function and a batch normalization after the first layer, and a Softplus activation function after the second layer, which produces the non-negative **F** affiliation weights.

Our method is based on the Bernoulli-Poisson graph generative model (20, 30, 31), in which the entries of the adjacency matrix *A*_*u ν*_ are sampled according to a Bernoulli distribution parameterized by the community affiliations:

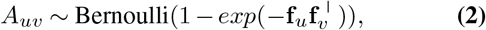

where **f**_*u*_ = *F*_*u*:_ and **f**_*ν*_ = *F*_*ν*:_ are the row vectors of the community affiliation matrix **F** of nodes *u* and *ν*, respectively. Further details behind the rationale of this model can be found in the Supplementary Material.

We approximate the negative log-likelihood of the Bernoulli-Poisson model by the following formula, which defines the Bernoulli-Poisson loss:

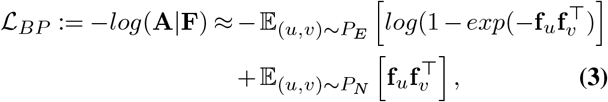

where *P*_*E*_ and *P*_*N*_ represent uniform distributions over edges and non-edges in the graph, respectively. These distributions are obtained through structured negative sampling. Specifically, for every positive edge sampled, a corresponding negative edge is also sampled, ensuring that the starting node is the same for both the positive and negative edges.

### Refining the identified modules with known biological pathways via constrained logistic regression

The second component begins by converting the non-negative module affiliations into probabilities, denoted as 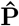 which represent the probability of a gene belonging to a module. This conversion is achieved by applying a scaled hyperbolic tangent function element-wise to the community affiliation matrix **F :**

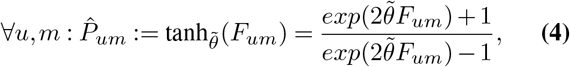

where 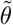 is a shared learnable scaling parameter.

Following this, for each *k* pathway database (*k ∈* {1, …, *L* }, where the *k* pathway database contains *K*^(*k*)^ number of pathways), individual linear layers are applied in parallel to the 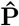 module probabilities, each employing a sigmoid activation function:

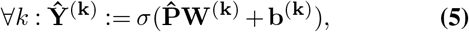

where Ŷ ^(**k**)^ are the *N* × *K*^(*k*)^ dimensional predicted outputs of the linear layers, and **W**^(**k**)^ and **b**^(**k**)^ are the *k*-th parallel linear layer’s weight matrix and bias vector, respectively. This step basically performs a binary classification over each individual pathway in each pathway database (i.e. a multilabel classification for each pathway database), where we denote the ground truth annotations as 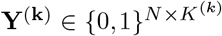in the *k* pathway database, i.e., 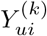 equals 1 if gene *u* is annotated with pathway *i* in the *k* pathway database and 0 otherwise.

We use a binary cross entropy loss to measure the difference between the predictions and the ground truth, and compute their sum for all *L* pathway databases and for each pathway, i.e.,

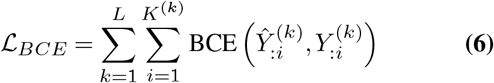

Note that the weights of the final parallel linear layers are constrained to be non-negative, denoted as 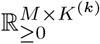, whereas the **b**^(**k**)^ biases of these layers are not subject to this constraint. Essentially, these final parallel layers function as *multivariable logistic regressions*, providing a coherent interpretation. Namely, the weight 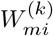 quantifies the increase in log odds of a gene being part of the *i*-th pathway in the *k* pathway database, given that the gene belongs to the *m* module. Formally, for gene *u* and the *i*-th pathway in the *k* pathway database:

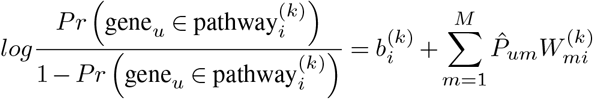

A notable advantage of this formulation is the comparability of the 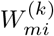 weight values. For each module *m*, the de-scending order of 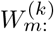 values directly corresponds to the relevance of pathways to the module. Conversely, for each pathway *i*, the descending order of 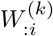 weights mirrors the significance of modules to the pathway’s function.

The non-negativity constraint on the weight matrices in the final layers has two purposes: Firstly, this deliberate reduction in the model’s representational capacity serves as a robust regularization technique. Secondly, this formulation is rooted in our 2nd assumption that functional modules are organized into a complex, hierarchical system that encompasses known biological pathways. This constraint reduces the model’s representational capacity to capture only simple additive interactions among modules, disallowing any complex combinations. This formalization can also be seen as a hierarchical model, where modules share a common weight of their genes being annotated with a pathway (i.e. the bias) while also having individual weights. As a result, the model ascertains a positive weight for certain, potentially overlapping modules relevant to a specific pathway, and it acquires a high negative bias value, rendering all other modules irrelevant to that pathway.

The objective function minimized during the end-to-end learning of the entire model comprises two parts: the Bernoulli-Poisson loss and the sum of all binary crossentropy losses of all pathways:

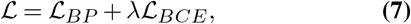

where *λ* is a hyper-parameter of the model.

#### Implementation details

We implemented our method using PyTorch v2.0.1 and PyTorch Geometric v2.3.1.

For each epoch during training, we shuffle all positive edges in the network, divide them into ten equal parts, and perform a training step for each part. We randomly split the pathway databases, used as model outputs, into training (80%) and validation (20%) pathways. During training, we use all annotated genes for the training pathway set, while for validation pathways, we use a subset of 80% randomly selected genes. We evaluate performance for each pathway database based on the excluded 20% genes from the validation pathway annotations, using the F1 score. We trained the model for 5000 epochs, selecting the model that achieved the highest mean F1 score across all pathway databases on the validation pathways.

We identified modules with varying module sizes, denoted by *M* (the dimensions of the hidden representations of **F**), specifically using module sizes of 500, 600, 700, 800, 900, and 1000. Our model automatically assigns genes to the identified modules by optimizing the 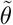 scaling parameter in Eq. 4 while we convert community affiliation strengths into module assignment probabilities. We still face a question regarding the appropriate probability cutoff for module assignment. We opt for a natural cutoff of 0.5, meaning we assign a gene to a module if the probability of its belonging to the module surpasses this threshold. Further details on finding the optimal settings of other hyper-parameters can be found in the Supplementary Methods.

We employed the AdamW optimizer with a starting learning rate of 0.001 and a Step learning rate scheduler, which decays the learning rate for each parameter group by 0.85 every 250 epochs. We performed all experiments on a Tesla V100 GPU.

#### Evaluation

Our evaluation method follows a similar framework to the Disease Module Identification DREAM Challenge, a community effort addressing the problem of disease module identification in complex molecular networks (32). To assess the model’s efficacy in identifying biologically relevant modules, we empirically examined the modules based on their association with complex traits and diseases, leveraging three distinct data sources and employing three different computational methods across them to mitigate biases in the assessment. Details of the data sources and evaluation methods are as follows:

#### GWAS Atlas

We used gene-level genome-wide association data from the GWAS Atlas project Release 3 (26), specifically the 1211 UK Biobank-specific gene-level summary statistics computed by the MAGMA software. Genes were ranked based on their p-values, and for each identified module, a Gene Set Enrichment Analysis using the `fgsea` R package (33) was performed to assess the enrichment of module genes in the rankings.

#### FinnGen

We used GWAS data from the FinnGen project (version DF9, Kurki et al. (34)), selecting the top 100 traits by prevalence that could be mapped to level 3 ICD-10 codes. Gene-level summary statistics were computed using the Pascal tool (35). Subsequently, for each module, an aggregated p-value was computed with Pascal, which employs a modified Fisher method to determine enrichment in high-scoring genes.

#### DisGeNET

We used all curated gene-disease associations from the DisGeNET database (version 7.0, Piñero et al. (36)) and selected diseases with at least 10 associated genes in the network. This resulted in 956 diseases associated with 6326 unique genes. An over-representation analysis was conducted based on a standard hypergeometric statistical test to evaluate the significance of the modules.

Two composite scores were derived to assess a method’s ability to identify biologically meaningful modules:

1.) The proportion of traits in each data source for which at least one significant module was identified (assessing the method’s ability to discover at least one disease mechanism for a particular disease).

2.) The proportion of modules in each data source for which at least one significant disease/trait was identified (evaluating the method’s capability to ascertain a relevant context for a module in which it is biologically meaningful).

The total score was computed as the summation of the scores from each evaluation data source, with a 5% false discovery rate (FDR) employed as the significance threshold.

#### Baseline methods

We chose several widely used and state-of-the-art community detection algorithms as baseline comparisons for GNN4DM. Utilizing the `cdlib` python package (37), we identified overlapping and non-overlapping modules employing various commonly used methods (see Supplementary Table S1). Additionally, we employed the top three high-performing state-of-the-art algorithms from the DREAM challenge, which focused on predicting non-overlapping modules ranging from 3 to 100 nodes. We also utilized the NOCD and UCoDe GNN-based methods to determine overlapping communities of the genes in the network. For these algorithms, the same input features were used as in the case of our method.

For all methods, we used the same underlying network derived from the STRING database. We performed an extensive grid search for algorithms with hyper-parameters to determine the optimal hyper-parameterization (see Supplementary Table S1), measured by the highest modularity density score (38). For NOCD and UCoDe, we specified the desired number of modules to match that of our method.

Besides, we also evaluated the pathway databases utilized for fine-tuning the module representations and the three subontologies of the Gene Ontology, treating them as results from a hypothetical overlapping module identification algorithm.

## Results

### Identification of modules in the human interactome

We applied the GNN4DM framework on the human interactome derived from the STRING protein-protein interaction database to identify its overlapping functional modules. The primary variable parameter in our method was the maximum module count, for which we experimented with a range from 500 to 1000 in increments of 100. To evaluate the concordance across these values, we quantified all pairwise overlaps between the various sets of modules by categorizing them as strong, sub-module, or partial (see Supplementary Methods). Our analysis indicated a substantial agreement among the chosen module counts, predominantly characterized by a high percentage of partial overlaps (see Supplementary Figure S1). Besides, increasing the maximum module count parameter tended to yield smaller sub-modules nested within those identified at lower counts. These observations, together with the generally low percentage of strong overlaps, suggest that the granularity of the modules varies *continuously* by increasing the maximum module count. This is also evident by the inverse relationship between the average module size and the maximum module count (see Supplementary Table S2). We also applied several baseline methods to identify both overlapping and non-overlapping modules within the network. Comprehensive descriptive statistics for all these methods, alongside the pathway databases used for training GNN4DM, and the Gene Ontology database, are detailed in Supplementary Table S2. The number and average size of modules identified varied significantly across methods, with module counts ranging from 8 (Eigenvector) to 10965 (Graph Entropy).

GNN4DM and the other two GNN-based methods showed notable differences. NOCD consistently generated far fewer modules than the pre-set maximum, with module counts ranging between 14 to 16. Both NOCD and UCoDe yielded modules with considerably large average sizes (1152.0-1283.6 for NOCD, 1359.2-1611.5 for UCoDe) but demonstrated low coverage (0.85-0.86 for NOCD, 0.74-0.84 for UCoDe). GNN4DM, on the other hand, consistently produced a module count close to its set maximum, with average module sizes between 72.3 and 120.8, and achieved nearly complete coverage (0.99).

As GNN4DM utilizes various pathway databases to refine its internal representations corresponding to the modules, we investigated whether its identified modules simply replicate these pathways or reveal additional information. Our analysis indicated that while GNN4DM aligns with pathway databases, especially with Reactome, it exhibits closer similarity to other methods such as the DREAM methods, CPM, and Graph Entropy, suggesting that GNN4DM identifies modules that are distinct from its training pathways (see Supplementary Results).

### Evaluation of functional disease modules

The evaluation of predicted modules poses a challenge due to the absence of a ground truth for ’correct’ modules against which the identified modules can be compared. To assess the capability of GNN4DM and other methods in identifying biologically relevant *functional disease modules*, we analyzed their associations with complex traits and diseases. This involved using GWAS data from GWAS Atlas and FinnGen, and curated gene-disease associations from DisGeNET. For subsequent analyses, we limited each method’s identified modules to 2-1000 genes. This restriction was based on the assumption that modules exceeding 1000 genes may lack biological specificity, potentially representing broad cellular functions instead of distinct disease-related processes, or could result from the method’s inability to cluster a large group of genes into smaller sub-modules (see Supplementary Table S3 for descriptive statistics after filtering).

First, we calculated a total score for each method, defined as the cumulative proportion of traits for which at least one significant module was identified. Our method, GNN4DM, outperformed all other methods in this regard, regardless of the maximum module count set within it (see Figure 2A). The highest score was observed with 1000 modules (total score = 2.47), surpassing even the scores of pathway databases used in training (ranging from 1.02 to 2.19). This score was also higher than, or on par with, the score achieved by Gene Ontology (2.46), which wasn’t part of our training set. Specifically, GNN4DM with 1000 modules identified significant modules for the highest proportion of traits in the GWAS Atlas (62.5%), exceeding the next best method (Node Perception, 53.1%) by 9.4 percentage points, and in FinnGen (90%), outperforming the second-best (Core Expansion, 86%) by 4 percentage points. For DisGeNET, GNN4DM identified at least one significant module for 94.6% of diseases, closely matching the top-performing method (Core Expansion, 95.7%).

**Fig. 2.**
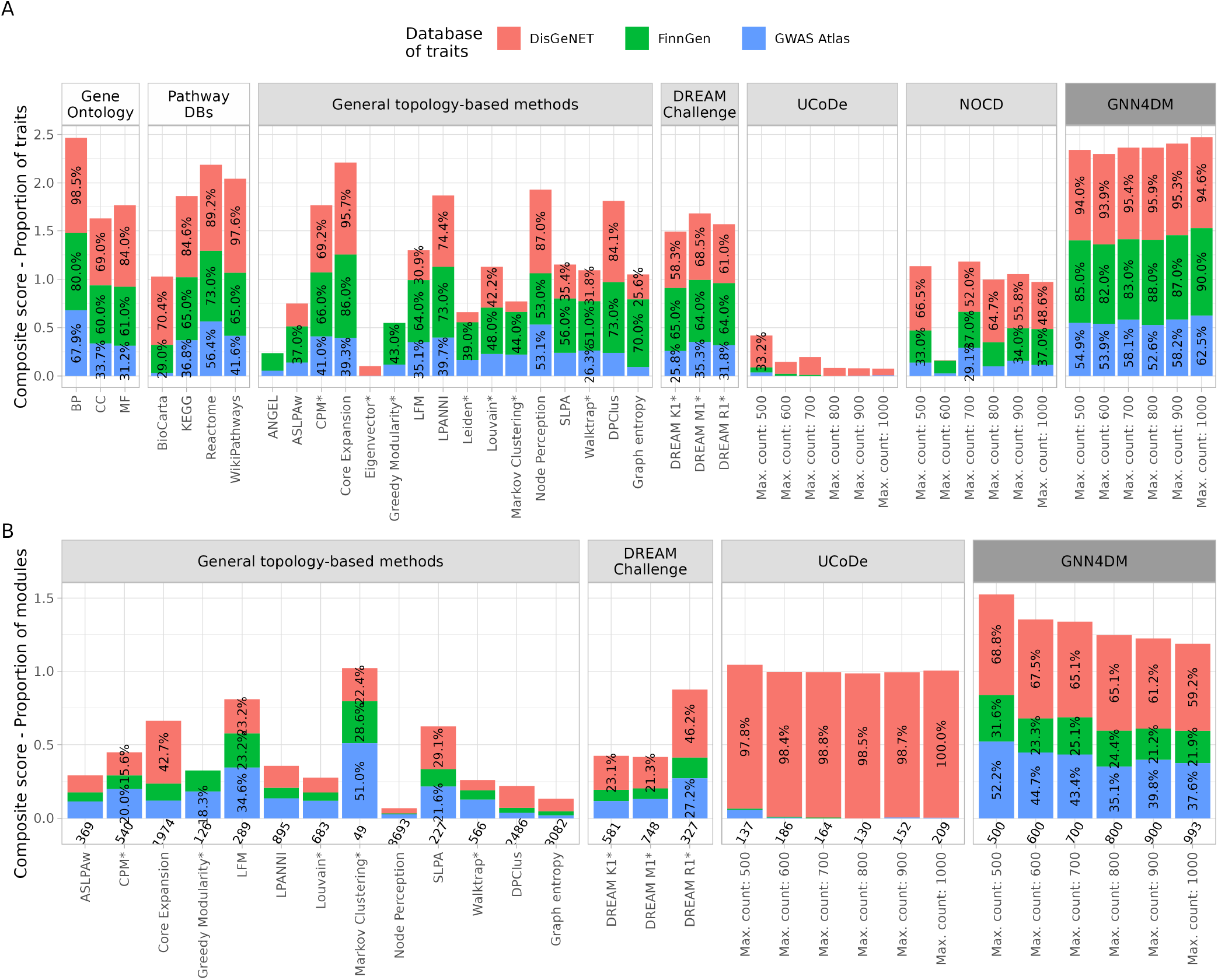
Performance evaluation of module identification methods and pathway databases. (**A**) Composite scores defined by the cumulative proportion of traits for which at least one significant module was identified across each data source. (**B**) Composite scores defined by the cumulative proportion of modules for which at least one significant trait was identified across each data source. Values under the columns indicate the number of modules in the size range of 2 to 1000. For part B, pathway databases and methods that identified fewer than 25 modules are omitted. Column groups with a white background header color indicate biological databases (in A), light gray indicates baseline methods, and dark gray indicates GNN4DM (on A&B). Percentage values inside the columns show the proportion of traits (in A), or modules (in B) that are significantly enriched/over-represented in the corresponding trait data source. Only percentages greater than 25% (on A) or 15% (on B) are shown. Method names denoted with an asterisk correspond to non-overlapping module identification methods.

Next, we computed another total score for each method, defined as the cumulative proportion of modules associated with at least one disease or trait. This score quantifies a method’s ability to discover a relevant context for their modules in which these are biologically meaningful. For this analysis, we omitted pathway databases to avoid selection bias, and methods that identified fewer than 25 modules to avoid bias due to low module counts. Again, GNN4DM consistently surpassed all other methods, regardless of its maximum module count setting (see Figure 2B), demonstrating a high proportion of significant modules across all three trait datasets.

The same qualitative trends hold when using more stringent significance thresholds (see Supplementary Figures S2 and S3). Overall, these findings underscore GNN4DM’s effectiveness in detecting a relatively high proportion of modules associated with a broad spectrum of diseases and complex traits, highlighting its strong capability in functional disease module identification.

### Case studies - Interpretation of highly enriched disease modules

We selected and visualized two highly enriched modules to demonstrate the interpretation of modules identified by GNN4DM, highlighting their associated diseases and relevant pathways.

#### Module 1

A notably enriched multimorbidity module (see Supplementary Figures S4A), derived using a maximum module count of 500, showed significant enrichment across the trait datasets: 39% of the diseases in the FinnGen dataset (39 out of the 100 most prevalent diseases with 3-level ICD codes), 11.8% in GWAS Atlas traits (143 out of 1211), and in a lesser extent, 2.82% in DisGeNET disorders (27 out of 956). This module is composed of four highly interconnected sub-parts, containing: 1) histone-coding genes, including core histones (H2A, H2B, H3, H4) and the H1 linker histone, crucial for DNA packaging and gene expression regulation. Also, in their extranuclear form, these histones are known to enhance host defense functions and contribute to inflammatory responses (39); 2) the Human Leukocyte Antigen complex, corresponding to MHC class II (DP, DM, DO, DQ, DR), which plays an essential role in the immune response against extracellular pathogens; 3) the CENP-A NAC/CAD kinetochore complex, vital for chromosome segregation during cell division, thereby maintaining genetic stability; and 4) cytokine-coding genes, involved in Th1 and Th2 immune responses, acting as key modulators of the body’s immune reaction.

To further understand the module’s biological relevance, we inspected the weights GNN4DM learned for the module in association with the known biological pathways. The module’s strong relation to immune-related pathways, including Th1/Th2, CTLA4, IL-4, and NF-kB signaling, emphasizes its role in immune regulation. Additionally, its association with pathways related to chromosome maintenance, gene expression regulation, and cellular senescence highlights its broader involvement in genetic stability and cellular aging processes (see Supplementary Figures S4C). This is also reflected by the module’s disease associations (see Supplementary Figures S4B) with various immunological disorders (like type 1 diabetes, asthma, celiac disease, allergic rhinitis, and eczema) and conditions tied to genomic instability (such as malignancies and schizophrenia).

#### Module 2

Another module highly over-represented in 17.4% among DisGeNET’s disorders (166 out of 956) is shown in Figure 3A. This module is comprised of a single densely interconnected component, primarily formed by cytokines, including chemokines from the CC and CXC sub-families, several interleukins, cytokine receptors, various regulatory protein-coding genes, and several other genes. The genes within the module are predominantly involved in immune response processes, inflammation, and r elated pathomechanisms. Notably, it encompasses both pro-inflammatory (e.g. *IL1B, IL18*) and anti-inflammatory (e.g. *IL10, IL4*) cytokine-coding genes, as well as interleukins with dual roles (e.g. *IL6*), suggesting a function in regulating the balance between these contrasting immune processes.

**Fig. 3.**
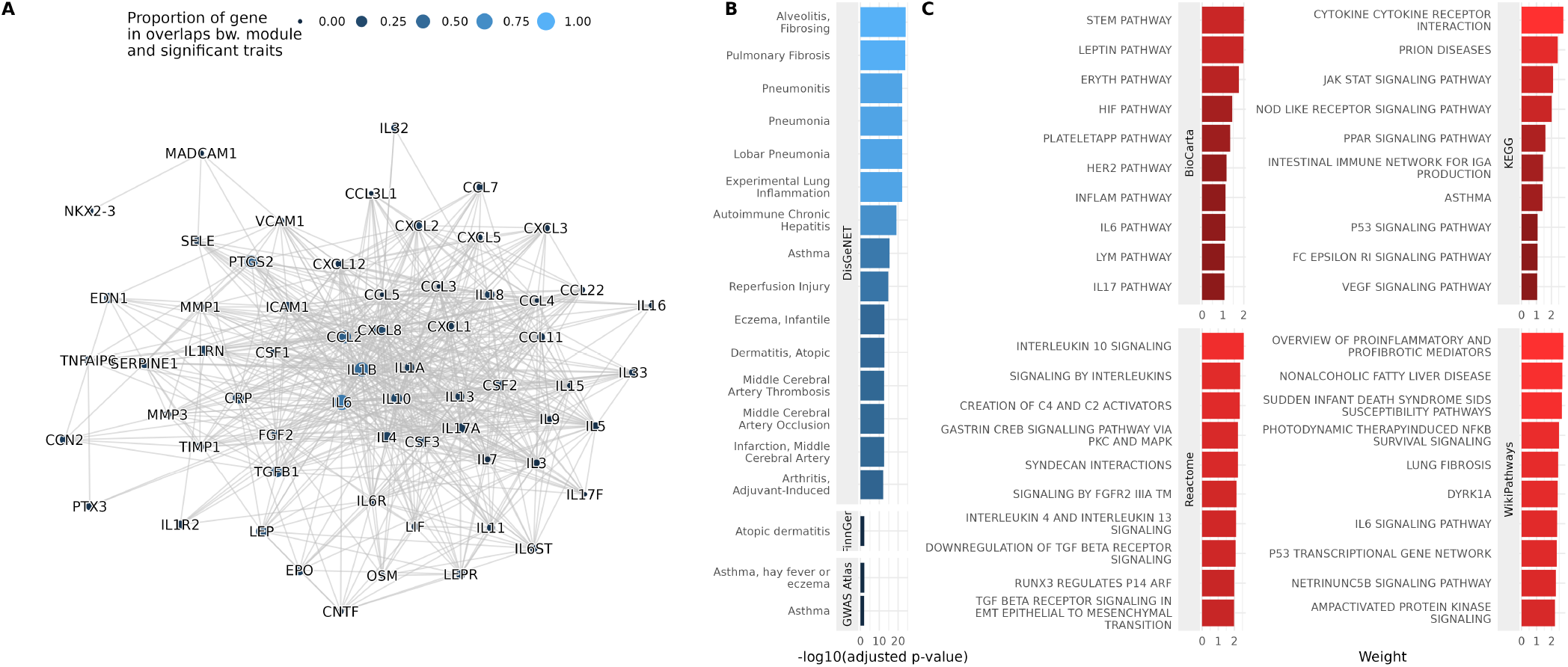
A highly enriched multimorbidity module. (**A**) The graph displays a subnetwork of the STRING PPI network, representing the module. The size and color of each node denote the gene’s proportion among disease-associated genes in those diseases that show significant overlap with the module, as determined by DisGeNET. (**B**) The top (at most) 15 diseases significantly associated with the module, as identified in the three evaluation datasets. (**C**) The top 10 pathways in each pathway database that are relevant to the module. The bars show the weight GNN4DM learned for the module in association with the corresponding pathway.

The potential functions of the module are also reflected in its related pathways, which include the Cytokines and inflammatory response pathway, the Cytokine-cytokine receptor interaction pathway, and several interleukin signaling pathways, among others (see Figure 3C). The associated diseases include several lung diseases, such as pneumonia, pulmonary fibrosis, and asthma; autoimmune disorders, such as rheumatoid arthritis and autoimmune chronic hepatitis; inflammatory diseases, such as atopic dermatitis and eczema; and various others (see Figure 3B, and Supplementary Figure S5).

## Conclusion

In this study, we introduced GNN4DM, a novel graph neural network-based approach for identifying functional disease modules in protein-protein interaction networks. GNN4DM has demonstrated strong capability in detecting biologically meaningful modules by integrating a substantial amount of genomic data and aligning the modules with known biological pathways in an end-to-end manner. Our method not only outperforms existing techniques in module identification but also provides valuable interpretations by linking modules to specific pathways. GNN4DM’s case studies further illustrate its utility in revealing complex disease mechanisms. Overall, GNN4DM represents a step forward to understanding complex diseases, providing a new tool for analyzing their molecular underpinnings.

## Author contributions statement

A.G. and P.A. conceived the experiments. A.G. conducted the experiments and analyzed the results. A.G. and P.A. wrote and reviewed the manuscript.

## Acknowledgments

We want to acknowledge the participants and investigators of the FinnGen study. AI-assisted tools were used only for the purpose of correcting written text.

## References

1. Leland H. Hartwell, John J. Hopfield, Stanislas Leibler, and Andrew W. Murray. From molecular to modular cell biology. Nature, 402:C47–C52, 12 1999. ISSN 0028-0836. doi: 10.1038/35011540.

2. Koyel Mitra, Anne-Ruxandra Carvunis, Sanath Kumar Ramesh, and Trey Ideker. Integrative approaches for finding modular structure in biological networks. Nature Reviews Genetics, 14:719–732, 10 2013. ISSN 1471-0056. doi: 10.1038/nrg3552.

3. Roded Sharan, Igor Ulitsky, and Ron Shamir. Network-based prediction of protein function. Molecular Systems Biology, 3, 1 2007. ISSN 1744-4292. doi: 10.1038/msb4100129.

4. M Oti and HG Brunner. The modular nature of genetic diseases. Clinical Genetics, 71(1): 1–11, 2007. doi: 10.1111/j.1399-0004.2006.00708.x.

5. Jörg Menche, Amitabh Sharma, Maksim Kitsak, Susan Dina Ghiassian, Marc Vidal, Joseph Loscalzo, and Albert-László Barabási. Uncovering disease-disease relationships through the incomplete interactome. Science, 347(6224):1257601, 2015. doi: 10.1126/science.1257601.

6. Monica Chagoyen and Florencio Pazos. Characterization of clinical signs in the human interactome. Bioinformatics, 32(12):1761–1765, 02 2016. ISSN 1367-4803. doi: 10.1093/bioinformatics/btw054.

7. Joseph Loscalzo and Albert-Laszlo Barabasi. Systems biology and the future of medicine. WIREs Systems Biology and Medicine, 3(6):619–627, 2011. doi: 10.1002/wsbm.144.

8. Hazel Nicolette Manners, Swarup Roy, and Jugal K. Kalita. Intrinsic-overlapping co-expression module detection with application to alzheimer’s disease. Computational Biology and Chemistry, 77:373–389, 12 2018. ISSN 14769271. doi: 10.1016/j.compbiolchem.2018.10.014.

9. Tao Wang, Qidi Peng, Bo Liu, Yongzhuang Liu, and Yadong Wang. Disease module identification based on representation learning of complex networks integrated from gwas, eqtl summaries, and human interactome. Frontiers in Bioengineering and Biotechnology, 8, 5 2020. ISSN 2296-4185. doi: 10.3389/fbioe.2020.00418.

10. Shiwen Zhao and Shao Li. A co-module approach for elucidating drug–disease associations and revealing their molecular basis. Bioinformatics, 28:955–961, 4 2012. ISSN 1367-4811. doi: 10.1093/bioinformatics/bts057.

11. Peng Ni, Jianxin Wang, Ping Zhong, Yaohang Li, Fang-Xiang Wu, and Yi Pan. Constructing disease similarity networks based on disease module theory. IEEE/ACM Transactions on Computational Biology and Bioinformatics, 17:906–915, 5 2020. ISSN 1545-5963. doi:10.1109/TCBB.2018.2817624.

12. Gergely Palla, Imre Derényi, Illés Farkas, and Tamás Vicsek. Uncovering the overlapping community structure of complex networks in nature and society. Nature, 435:814–818, 6 2005. ISSN 0028-0836. doi: 10.1038/nature03607.

13. Anne-Claude Gavin, Markus Bösche, Roland Krause, Paola Grandi, Martina Marzioch, Andreas Bauer, Jörg Schultz, Jens M. Rick, Anne-Marie Michon, Cristina-Maria Cruciat, Marita Remor, Christian Höfert, Malgorzata Schelder, Miro Brajenovic, Heinz Ruffner, Alejandro Merino, Karin Klein, Manuela Hudak, David Dickson, Tatjana Rudi, Volker Gnau, Angela Bauch, Sonja Bastuck, Bettina Huhse, Christina Leutwein, Marie-Anne Heurtier, Richard R. Copley, Angela Edelmann, Erich Querfurth, Vladimir Rybin, Gerard Drewes, Manfred Raida, Tewis Bouwmeester, Peer Bork, Bertrand Seraphin, Bernhard Kuster, Gitte Neubauer, and Giulio Superti-Furga. Functional organization of the yeast proteome by systematic analysis of protein complexes. Nature, 415:141–147, 1 2002. ISSN 0028-0836. doi: 10.1038/415141a.

14. E. Ravasz, A. L. Somera, D. A. Mongru, Z. N. Oltvai, and A.-L. Barabási. Hierarchical organization of modularity in metabolic networks. Science, 297(5586):1551–1555, 2002. doi: 10.1126/science.1073374.

15. Orkun S Soyer. Emergence and maintenance of functional modules in signaling pathways. BMC Evolutionary Biology, 7:205, 2007. ISSN 1471-2148. doi: 10.1186/1471-2148-7-205.

16. Jeff Clune, Jean-Baptiste Mouret, and Hod Lipson. The evolutionary origins of modularity. Proceedings of the Royal Society B: Biological Sciences, 280:20122863, 3 2013. ISSN 0962-8452. doi: 10.1098/rspb.2012.2863.

17. Henok Mengistu, Joost Huizinga, Jean-Baptiste Mouret, and Jeff Clune. The evolutionary origins of hierarchy. PLOS Computational Biology, 12:e1004829, 6 2016. ISSN 1553-7358. doi: 10.1371/journal.pcbi.1004829.

18. William L. Hatleberg and Veronica F. Hinman. Chapter two - modularity and hierarchy in biological systems: Using gene regulatory networks to understand evolutionary change. In Scott F. Gilbert, editor, Evolutionary Developmental Biology, volume 141 of Current Topics in Developmental Biology, pages 39–73. Academic Press, 2021. doi: 10.1016/bs.ctdb.2020.11.004.

19. Ichcha Manipur, Maurizio Giordano, Marina Piccirillo, Seetharaman Parashuraman, and Lucia Maddalena. Community detection in protein-protein interaction networks and applications. IEEE/ACM Transactions on Computational Biology and Bioinformatics, 20:217–237, 1 2023. ISSN 1545-5963. doi: 10.1109/TCBB.2021.3138142.

20. Oleksandr Shchur and Stephan Günnemann. Overlapping community detection with graph neural networks, 2019.

21. Xiao Ye, Yulin Wu, Jiangsheng Pi, Hong Li, Bo Liu, Yadong Wang, and Junyi Li. Deepgmd: A graph-neural-network-based method to detect gene regulator module. IEEE/ACM transactions on computational biology and bioinformatics, 19:3366–3373, 2022. ISSN 1557-9964. doi: 10.1109/TCBB.2021.3114281.

22. Atefeh Moradan, Andrew Draganov, Davide Mottin, and Ira Assent. Ucode: unified community detection with graph convolutional networks. Machine Learning, 112:5057–5080, 12 2023. ISSN 0885-6125. doi: 10.1007/s10994-023-06402-0.

23. Damian Szklarczyk, Annika L. Gable, David Lyon, Alexander Junge, Stefan Wyder, Jaime Huerta-Cepas, Milan Simonovic, Nadezhda T. Doncheva, John H. Morris, Peer Bork, Lars J. Jensen, and Christian Von Mering. String v11: protein-protein association networks with increased coverage, supporting functional discovery in genome-wide experimental datasets. Nucleic acids research, 47:D607–D613, 1 2019. ISSN 1362-4962. doi: 10.1093/NAR/GKY1131.

24. Justin K. Huang, Daniel E. Carlin, Michael Ku Yu, Wei Zhang, Jason F. Kreisberg, Pablo Tamayo, and Trey Ideker. Systematic evaluation of molecular networks for discovery of disease genes. Cell Systems, 6:484–495.e5, 4 2018. ISSN 24054712. doi: 10.1016/j.cels.2018.03.001.

25. François Aguet, Shankara Anand, Kristin G. Ardlie, Stacey Gabriel, Gad A. Getz, Aaron Graubert, Kane Hadley, Robert E. Handsaker, Katherine H. Huang, Seva Kashin, Xiao Li, Daniel G. MacArthur, Samuel R. Meier, Jared L. Nedzel, Duyen T. Nguyen, Ayellet V. Segrè, Ellen Todres, Brunilda Balliu, Alvaro N. Barbeira, Alexis Battle, Rodrigo Bonazzola, Andrew Brown, Christopher D. Brown, Stephane E. Castel, Donald F. Conrad, Daniel J. Cotter, Nancy Cox, Sayantan Das, Olivia M. de Goede, Emmanouil T. Dermitzakis, Jonah Einson, Barbara E. Engelhardt, Eleazar Eskin, Tiffany Y. Eulalio, Nicole M. Ferraro, Elise D. Flynn, Laure Fresard, Eric R. Gamazon, Diego Garrido-Martín, Nicole R. Gay, Michael J. Gloudemans, Roderic Guigó, Andrew R. Hame, Yuan He, Paul J. Hoffman, Farhad Hormoz- diari, Lei Hou, Hae Kyung Im, Brian Jo, Silva Kasela, Manolis Kellis, Sarah Kim-Hellmuth, Alan Kwong, Tuuli Lappalainen, Xin Li, Yanyu Liang, Serghei Mangul, Pejman Mohammadi, Stephen B. Montgomery, Manuel Muñoz-Aguirre, Daniel C. Nachun, Andrew B. Nobel, Meritxell Oliva, YoSon Park, Yongjin Park, Princy Parsana, Abhiram S. Rao, Ferran Reverter, John M. Rouhana, Chiara Sabatti, Ashis Saha, Matthew Stephens, Barbara E. Stranger, Benjamin J. Strober, Nicole A. Teran, Ana Viñuela, Gao Wang, Xiaoquan Wen, Fred Wright, Valentin Wucher, Yuxin Zou, Pedro G. Ferreira, Gen Li, Marta Melé, Esti Yeger-Lotem, Mary E. Barcus, Debra Bradbury, Tanya Krubit, Jeffrey A. McLean, Liqun Qi, Karna Robinson, Nancy V. Roche, Anna M. Smith, Leslie Sobin, David E. Tabor, Anita Undale, Jason Bridge, Lori E. Brigham, Barbara A. Foster, Bryan M. Gillard, Richard Hasz, Marcus Hunter, Christopher Johns, Mark Johnson, Ellen Karasik, Gene Kopen, William F. Leinweber, Alisa McDonald, Michael T. Moser, Kevin Myer, Kimberley D. Ramsey, Brian Roe, Saboor Shad, Jeffrey A. Thomas, Gary Walters, Michael Washington, Joseph Wheeler, Scott D. Jewell, Daniel C. Rohrer, Dana R. Valley, David A. Davis, Deborah C. Mash, Philip A. Branton, Laura K. Barker, Heather M. Gardiner, Maghboeba Mosavel, Laura A. Siminoff, Paul Flicek, Maximilian Haeussler, Thomas Juettemann, W. James Kent, Christopher M. Lee, Conner C. Powell, Kate R. Rosenbloom, Magali Ruffier, Dan Sheppard, Kieron Taylor, Stephen J. Trevanion, Daniel R. Zerbino, Nathan S. Abell, Joshua Akey, Lin Chen, Kathryn Demanelis, Jennifer A. Doherty, Andrew P. Feinberg, Kasper D. Hansen, Peter F. Hickey, Farzana Jasmine, Lihua Jiang, Rajinder Kaul, Muhammad G. Kibriya, Jin Billy Li, Qin Li, Shin Lin, Sandra E. Linder, Brandon L. Pierce, Lindsay F. Rizzardi, Andrew D. Skol, Kevin S. Smith, Michael Snyder, John Stamatoyannopoulos, Hua Tang, Meng Wang, Latarsha J. Carithers, Ping Guan, Susan E. Koester, A. Roger Little, Helen M. Moore, Concepcion R. Nierras, Abhi K. Rao, Jimmie B. Vaught, and Simona Volpi. The gtex consortium atlas of genetic regulatory effects across human tissues. Science, 369:1318–1330, 9 2020. ISSN 0036-8075. doi: 10.1126/science.aaz1776.

26. Kyoko Watanabe, Sven Stringer, Oleksandr Frei, Maša Umićević Mirkov, Christiaan de Leeuw, Tinca J C Polderman, Sophie van der Sluis, Ole A Andreassen, Benjamin M Neale, and Danielle Posthuma. A global overview of pleiotropy and genetic architeture in complex traits. Nature genetics, 51:1339–1348, 9 2019. ISSN 1546-1718. doi: 10.1038/s41588-019-0481-0.

27. Aric A. Hagberg, Daniel A. Schult, and Pieter J. Swart. Exploring network structure, dynamics, and function using networkx. In Gaël Varoquaux, Travis Vaught, and Jarrod Millman, editors, Proceedings of the 7th Python in Science Conference, pages 11–15, Pasadena, CA USA, 2008.

28. Aravind Subramanian, Pablo Tamayo, Vamsi K. Mootha, Sayan Mukherjee, Benjamin L. Ebert, Michael A. Gillette, Amanda Paulovich, Scott L. Pomeroy, Todd R. Golub, Eric S. Lander, and Jill P. Mesirov. Gene set enrichment analysis: A knowledge-based approach for interpreting genome-wide expression profiles. Proceedings of the National Academy of Sciences, 102:15545–15550, 10 2005. ISSN 0027-8424. doi: 10.1073/pnas.0506580102.

29. Thomas N. Kipf and Max Welling. Semi-supervised classification with graph convolutional networks, 2017.

30. Ioannis Psorakis, Stephen Roberts, Mark Ebden, and Ben Sheldon. Overlapping community detection using bayesian non-negative matrix factorization. Phys. Rev. E, 83:066114, Jun 2011. doi: 10.1103/PhysRevE.83.066114.

31. Jaewon Yang and Jure Leskovec. Overlapping community detection at scale: A nonnegative matrix factorization approach. In Proceedings of the Sixth ACM International Conference on Web Search and Data Mining, WSDM ’13, page 587–596, New York, NY, USA, 2013. Association for Computing Machinery. ISBN 9781450318693. doi: 10.1145/2433396.2433471.

32. Sarvenaz Choobdar, Mehmet E. Ahsen, Jake Crawford, Mattia Tomasoni, Tao Fang, David Lamparter, Junyuan Lin, Benjamin Hescott, Xiaozhe Hu, Johnathan Mercer, Ted Natoli, Rajiv Narayan, Aravind Subramanian, Jitao D. Zhang, Gustavo Stolovitzky, Zoltán Kutalik, Kasper Lage, Donna K. Slonim, Julio Saez-Rodriguez, Lenore J. Cowen, Sven Bergmann, and Daniel Marbach. Assessment of network module identification across complex diseases. Nature Methods, 16:843–852, 9 2019. ISSN 1548-7091. doi: 10.1038/s41592-019-0509-5.

33. Alexey A. Sergushichev. An algorithm for fast preranked gene set enrichment analysis using cumulative statistic calculation. bioRxiv, 2016. doi: 10.1101/060012.

34. Mitja I. Kurki, Juha Karjalainen, Priit Palta, Timo P. Sipilä, Kati Kristiansson, Kati M. Donner, Mary P. Reeve, Hannele Laivuori, Mervi Aavikko, Mari A. Kaunisto, Anu Loukola, Elisa Lahtela, Hannele Mattsson, Päivi Laiho, Pietro Della Briotta Parolo, Arto A. Lehisto, Masahiro Kanai, Nina Mars, Joel Rämö, Tuomo Kiiskinen, Henrike O. Heyne, Kumar Veerapen, Sina Rüeger, Susanna Lemmelä, Wei Zhou, Sanni Ruotsalainen, Kalle Pärn, Tero Hiekkalinna, Sami Koskelainen, Teemu Paajanen, Vincent Llorens, Javier Gracia-Tabuenca, Harri Siirtola, Kadri Reis, Abdelrahman G. Elnahas, Benjamin Sun, Christopher N. Foley, Katriina Aalto-Setälä, Kaur Alasoo, Mikko Arvas, Kirsi Auro, Shameek Biswas, Argyro Bizaki-Vallaskangas, Olli Carpen, Chia-Yen Chen, Oluwaseun A. Dada, Zhihao Ding, Margaret G. Ehm, Kari Eklund, Martti Färkkilä, Hilary Finucane, Andrea Ganna, Awaisa Ghazal, Robert R. Graham, Eric M. Green, Antti Hakanen, Marco Hautalahti, Åsa K. Hedman, Mikko Hiltunen, Reetta Hinttala, Iiris Hovatta, Xinli Hu, Adriana Huertas-Vazquez, Laura Huilaja, Julie Hunkapiller, Howard Jacob, Jan-Nygaard Jensen, Heikki Joensuu, Sally John, Valtteri Julkunen, Marc Jung, Juhani Junttila, Kai Kaarniranta, Mika Kähönen, Risto Kajanne, Lila Kallio, Reetta Kälviäinen, Jaakko Kaprio, Nurlan Kerimov, Johannes Kettunen, Elina Kilpeläinen, Terhi Kilpi, Katherine Klinger, Veli-Matti Kosma, Teijo Kuopio, Venla Kurra, Triin Laisk, Jari Laukkanen, Nathan Lawless, Aoxing Liu, Simonne Longerich, Reedik Mägi, Johanna Mäkelä, Antti Mäkitie, Anders Malarstig, Arto Mannermaa, Joseph Maranville, Athena Matakidou, Tuomo Meretoja, Sahar V. Mozaffari, Mari E. K. Niemi, Marianna Niemi, Teemu Niiranen Christopher J. O’Donnell, Maén Obeidat, George Okafo, Hanna M. Ollila, Antti Palomäki, Tuula Palotie, Jukka Partanen, Dirk S. Paul, Margit Pelkonen, Rion K. Pendergrass, Slavé Petrovski, Anne Pitkäranta, Adam Platt, David Pulford, Eero Punkka, Pirkko Pussinen, Neha Raghavan, Fedik Rahimov, Deepak Rajpal, Nicole A. Renaud, Bridget Riley-Gillis, Rodosthenis Rodosthenous, Elmo Saarentaus, Aino Salminen, Eveliina Salminen, Veikko Salomaa, Johanna Schleutker, Raisa Serpi, Huei yi Shen, Richard Siegel, Kaisa Silander, Sanna Siltanen, Sirpa Soini, Hilkka Soininen, Jae Hoon Sul, Ioanna Tachmazidou, Kaisa Tasanen, Pentti Tienari, Sanna Toppila-Salmi, Taru Tukiainen, Tiinamaija Tuomi, Joni A. Turunen, Jacob C. Ulirsch, Felix Vaura, Petri Virolainen, Jeffrey Waring, Dawn Waerworth, Robert Yang, Mari Nelis, Anu Reigo, Andres Metspalu, Lili Milani, Tõnu Esko, Caroline Fox, Aki S. Havulinna, Markus Perola, Samuli Ripatti, Anu Jalanko, Tarja Laitinen, Tomi P. Mäkelä, Robert Plenge, Mark McCarthy, Heiko Runz, Mark J. Daly, and Aarno Palotie. Finngen provides genetic insights from a well-phenotyped isolated population. Nature, 613:508–518, 1 2023. ISSN 0028-0836. doi: 10.1038/s41586-022-05473-8.

35. David Lamparter, Daniel Marbach, Rico Rueedi, Zoltán Kutalik, and Sven Bergmann. Fast and rigorous computation of gene and pathway scores from snp-based summary statistics. PLOS Computational Biology, 12:e1004714, 1 2016. ISSN 1553-7358. doi: 10.1371/journal.pcbi.1004714.

36. Janet Piñero, Juan Manuel Ramírez-Anguita, Josep Saüch-Pitarch, Francesco Ronzano, Emilio Centeno, Ferran Sanz, and Laura I Furlong. The disgenet knowledge platform for disease genomics: 2019 update. Nucleic Acids Research, 11 2019. ISSN 0305-1048. doi: 10.1093/nar/gkz1021.

37. Giulio Rossetti, Letizia Milli, and Rémy Cazabet. The disgenet knowledge platform for diCdlib: a python library to extract, compare and evaluate communities from complex networks. Applied Network Science, 4:52, 12 2019. ISSN 2364-8228. doi: 10.1007/s41109-019-0165-9.

38. Shihua Zhang, Xue-Mei Ning, Chris Ding, and Xiang-Sun Zhang. The disgenet knowledge platform for diDetermining modular organization of protein interaction networks by maximizing modularity density. BMC Systems Biology, 4:S10, 9 2010. ISSN 1752-0509. doi: 10.1186/1752-0509-4-S2-S10.

39. Xia Li, Youyuan Ye, Kailan Peng, Zhuo Zeng, Li Chen, and Yanhua Zeng. Histones: The critical players in innate immunity. Frontiers in immunology, 13:1030610, 2022. ISSN 1664-3224. doi: 10.3389/fimmu.2022.1030610.

