## Supplementary Material for "GNN4DM: A Graph Neural Network-based method to identify overlapping functional disease modules"

András Gézsi, Péter Antal
Department of Measurement and Information Systems,
Budapest University of Technology and Economics, Budapest, Hungary



#### 1 Supplementary Results

##### 1.1 Similarity and complementarity of different module identification methods

As GNN4DM utilizes various pathway databases to refine its internal representations corresponding to the modules, a relevant question arises: do the modules identified by GNN4DM merely replicate these pathways, or do they capture extra layers of information? To address this, we conducted two analyses.

Initially, we measured all pairwise overlaps between GNN4DM-identified modules and pathways in the databases, classifying them as strong, sub-module, or partial overlaps (see Supplementary Methods). We observed a generally low rate of strong overlap, with less than 2% of GNN4DM modules closely matching any pathway from the databases. Conversely, partial overlaps were more common (see Supplementary Figure S6A). Meanwhile, the pathways often tend to be sub-modules of larger GNN4DM modules, suggesting that these modules represent compact, higher-level biological functions that encompass multiple known pathways (see Supplementary Figure S6B).

Subsequently, to discern whether different methods reveal similar or distinct modules, we calculated a pairwise similarity metric among module identification methods and visualized their relationships using multidimensional scaling (see Supplementary Figure S7 and Supplementary Methods) and a heatmap combined with hierarchical clustering (see Supplementary Figure S8). This analysis demonstrated that although GNN4DM variants show alignment with pathway databases, particularly Reactome, they exhibit closer similarity to other approaches, including DREAM methods, CPM, and Graph Entropy. This underscores that GNN4DM identifies modules distinct from the pathways used for training it.

#### 1.2 Enrichment of modules across the three trait datasets

To conduct a more in-depth analysis of modules identified by GNN4DM and their associations with traits, we generated pairwise scatter plots showing the proportion of traits associated with these modules in the trait datasets (see Supplementary Figure S9). A significant correlation is observed between GWAS Atlas and FinnGen, indicating that modules highly enriched in one database tend to be similarly enriched in the other (see Supplementary Figure S9A). However, the correlation between DisGeNET and the GWAS databases is markedly lower, or in some cases, even negative (see Supplementary Figures S9B and C). This discrepancy could be attributed to the differing disease prevalences across the datasets, particularly since DisGeNET includes a larger proportion of rare diseases. Additionally, the over-representation analysis used for DisGeNET tends to favor larger gene sets (as the mean number of associated genes is also larger in this dataset), unlike the enrichment methods applied to the other two databases.

### 2 Supplementary Methods

#### 2.1 Implementation details

We conducted exploratory experiments to find the optimal settings of other hyper-parameters, including the number of hidden GCN layers, the number of neurons, and the $\lambda$ hyper-parameter weighing the BCE loss component, among others. We also experimented with various graph neural network architectures, including Graph Attention layers and GraphSAGE, alongside different regularization techniques like dropout, and L1 or L2 regularization of the weight matrices. However, these alterations did not yield any noticeable improvements. Additionally, we attempted to incorporate the Gene Ontology database as target outputs for the model, yet this unexpectedly led to inferior results.

We show the finalized hyper-parameters for our model in Supplementary Table S4.

#### 2.2 A note on the Bernoulli-Poisson model

The rationale behind Eq. 2 in the main paper stems from our first assumption, namely: (1.a) Each gene $u$ belongs to a given functional module $m$ with a given strength $F_{um}$. The higher the weight is, the more likely that this gene interacts with other members of the same module, depending on their own weights. (1.b) A gene (or a gene product) may have various functional roles in the cell, which are carried out by different functional modules. This way, a gene may belong to multiple modules and, therefore, has multiple independent chances to interact with another gene $v$ if they both participate in multiple modules. In other words, the more modules two genes share, the higher the probability of their interaction (i.e., an edge appears in the graph). More formally, we assume that a pair of genes $(u, v)$ has a latent interaction of strength $S_{uv}^{(m)}$ within the $m$ module, and this strength variable has a Poisson distribution with mean $F_{um} \cdot F_{vm}$. Then, the aggregate interaction effect, denoted as $S_{uv}$, between nodes $u$ and $v$, is obtained by summing over all $m$ modules, namely $S_{uv} = \sum_m S_{uv}^{(m)}$. Then, according to the additivity rule of Poisson distributed random variables, $S_{uv} \sim \text{Pois}(\mathbf{f}_u \mathbf{f}_v^\top)$, where $\mathbf{f}_u = F_{u:}$ and $\mathbf{f}_v = F_{v:}$. The probability of the edge between $(u, v)$ amounts to the probability of $S_{uv}$ being non-zero, i.e.: $P(S_{uv} > 0) = 1 - P(S_{uv} = 0) = 1 - exp(-\mathbf{f}_u \mathbf{f}_v^\top)$, which is the same as Eq. 2.

#### 2.3 Similarity of module predictions

Adapting the approach from [3], we defined a similarity measure for comparing module predictions from different methods. We represented the set of modules from a method $m$ as a prediction vector $P_m$, with a length of $N(N-1)/2$, where $N$ represents the total number of genes in the network. Each vector element corresponds to a gene pair, denoting the count of modules encompassing both genes. To quantify the similarity between two methods $m1$ and $m2$, we computed their distance as:

$$D(m_1, m_2) = 1 - \frac{< P_{m1}, P_{m2} >}{||P_{m1}||_2 \cdot ||P_{m2}||_2},$$

where $< .,. >$ is the inner product and $||.||_2$ is the Euclidean norm. The resulting symmetric distance matrix $D$ served as input for multidimensional scaling (MDS) to reduce dimensionality in Figure S7. This distance matrix was also visualized as a heatmap in Supplementary Figure S8. This metric extends the approach of [3] for non-overlapping modules to overlapping ones.

#### 2.4 Overlap between trait-associated modules

In line with [3], we employed various metrics to quantify the overlap between modules from various methods. First, we evaluated the statistical significance of the overlaps using hypergeometric distribution. An overlap was deemed statistically significant if its Bonferroni-adjusted p-value was below 0.05, taking into account the total number of module pair comparisons conducted. Next, we quantified the overlap using the Jaccard index, which is the ratio of the size of the intersection to the size of the union of two modules (i.e., gene sets) $A$ and $B$:

$$J(A, B) = \frac{|A \cap B|}{|A \cup B|}$$

Additionally, to identify sub-modules, we also considered the proportion of genes in the first module present in the second module:

$$S(A, B) = \frac{|A \cap B|}{|A|}$$

Based on these metrics, we classified each statistically significant overlap between module $A$ and another module $B$ into three categories:

1. Strong overlap: $J(A, B) \geq 0.5$.
2. Sub-module: $J(A, B) < 0.5$ and $S(A, B) - J(A, B) \geq 0.5$.
3. Partial overlap: $J(A, B) < 0.5$ and $S(A, B) - J(A, B) < 0.5$.

We applied these classifications to get an overview of the granularity of different maximum module count settings in GNN4DM, and to compare the modules from GNN4DM with all pathways of the pathway databases utilized for training (see Supplementary Figures S1 and S6).

### 3 Supplementary Tables

Table S1: Hyper-parameterization and references of baseline methods.

| | Method | Overlaps | Hyper-parameters | Reference |
|---|---|---|---|---|
| General topology-based methods | CPM | ✗ | Resolution: from 0.01 to 0.1 by 0.01 | [15] |
| | Eigenvector | ✗ | None | [10] |
| | Greedy Modularity | ✗ | None | [5] |
| | Leiden | ✗ | None | [16] |
| | Louvain | ✗ | Resolution: from 0.1 to 1.0 by 0.1<br>Randomize: True | [2] |
| | Markov Clustering | ✗ | Inflation: from 1.1 to 1.8 by 0.1 | [6] |
| | Walktrap | ✗ | None | [11] |
| | ANGEL | ✓ | Threshold: from 0.1 to 1.0 by 0.1 | [12] |
| | ASLPAw | ✓ | None | [17] |
| | Core Expansion | ✓ | None | [4] |
| | DPClus | ✓ | Density th.: from 0.8 to 0.95 by 0.05<br>Property th.: from 0.3 to 0.7 by 0.1 | [1] |
| | Graph entropy | ✓ | None | [7] |
| | LFM | ✓ | Alpha: from 0.5 to 2.0 by 0.1 | [8] |
| | LPANNI | ✓ | Threshold: 0.01 | [9] |
| | Node Perception | ✓ | Threshold: from 0.2 to 0.8 by 0.2<br>Overlap th.: from 0.2 to 0.8 by 0.2 | [13] |
| | SLPA | ✓ | R: from 0.1 to 0.5 by 0.1 | [17] |
| DREAM Challenge | DREAM K1 | ✗ | None | [3, 14] |
| | DREAM M1 | ✗ | None | |
| | DREAM R1 | ✗ | None | |

Table S2: Descriptive statistics for the investigated module identification methods and pathway databases, unfiltered.

| | Method | Overlaps | Count of identified modules | Average size (range) | Node coverage | Mean number of modules per gene | Conductance | Modularity overlap |
|---|---|---|---|---|---|---|---|---|
| General topology-based methods | CPM | ✗ | 1583 | 10.0 (1 - 830) | 1.00 | 1.00 | 0.83 | 0.03 |
| | Eigenvector | ✗ | 8 | 1974.1 (845 - 3877) | 1.00 | 1.00 | 0.27 | 0.01 |
| | Greedy Modularity | ✗ | 130 | 121.5 (2 - 5611) | 1.00 | 1.00 | 0.35 | 0.27 |
| | Leiden | ✗ | 32 | 493.5 (5 - 1693) | 1.00 | 1.00 | 0.19 | 0.15 |
| | Louvain | ✗ | 686 | 23.0 (2 - 1577) | 1.00 | 1.00 | 0.47 | 0.13 |
| | Markov Clustering | ✗ | 52 | 303.7 (2 - 6990) | 1.00 | 1.00 | 0.41 | 0.10 |
| | Walktrap | ✗ | 899 | 17.6 (1 - 4656) | 1.00 | 1.00 | 0.64 | 0.13 |
| | ANGEL | ✓ | 17 | 762.1 (4 - 12790) | 0.82 | 1.00 | 0.16 | 0.51 |
| | ASLPAw | ✓ | 583 | 29.6 (1 - 6253) | 1.00 | 1.09 | 0.67 | 0.08 |
| | Core Expansion | ✓ | 1974 | 20.5 (2 - 276) | 0.82 | 3.12 | 0.81 | 0.06 |
| | DPClus | ✓ | 2486 | 6.0 (2 - 185) | 0.77 | 1.22 | 0.78 | -0.21 |
| | Graph entropy | ✓ | 10965 | 2.4 (1 - 198) | 1.00 | 1.66 | 0.95 | -0.07 |
| | LFM | ✓ | 291 | 100.2 (2 - 13988) | 1.00 | 1.85 | 0.44 | 0.07 |
| | LPANNI | ✓ | 899 | 24.2 (1 - 3233) | 1.00 | 1.38 | 0.56 | 0.15 |
| | Node Perception | ✓ | 8699 | 6.7 (2 - 2183) | 0.95 | 3.90 | 0.94 | -0.09 |
| | SLPA | ✓ | 267 | 55.6 (1 - 2523) | 0.94 | 1.00 | 0.53 | 0.09 |
| DREAM Challenge | DREAM K1 | ✗ | 581 | 27.2 (2 - 100) | 1.00 | 1.00 | 0.59 | 0.08 |
| | DREAM M1 | ✗ | 2029 | 7.8 (1 - 100) | 1.00 | 1.00 | 0.85 | -0.02 |
| | DREAM R1 | ✗ | 327 | 44.7 (3 - 93) | 0.93 | 1.00 | 0.61 | 0.04 |
| UCoDe | Max. count: 500 | ✓ | 498 | 1359.2 (5 - 2317) | 0.74 | 57.57 | 0.23 | 0.00 |
| | Max. count: 600 | ✓ | 599 | 1413.0 (12 - 2669) | 0.77 | 69.73 | 0.22 | 0.00 |
| | Max. count: 700 | ✓ | 698 | 1521.7 (5 - 2955) | 0.77 | 87.37 | 0.21 | 0.00 |
| | Max. count: 800 | ✓ | 800 | 1591.3 (2 - 2946) | 0.80 | 100.70 | 0.21 | 0.00 |
| | Max. count: 900 | ✓ | 899 | 1582.4 (9 - 3436) | 0.82 | 109.49 | 0.21 | 0.00 |
| | Max. count: 1000 | ✓ | 999 | 1611.5 (82 - 3287) | 0.84 | 121.14 | 0.22 | 0.00 |
| NOCD | Max. count: 500 | ✓ | 15 | 1199.7 (437 - 2240) | 0.85 | 1.33 | 0.35 | 0.02 |
| | Max. count: 600 | ✓ | 16 | 1152.0 (1 - 2226) | 0.86 | 1.36 | 0.39 | 0.01 |
| | Max. count: 700 | ✓ | 15 | 1215.5 (352 - 2315) | 0.86 | 1.34 | 0.34 | 0.02 |
| | Max. count: 800 | ✓ | 14 | 1277.8 (13 - 2231) | 0.85 | 1.33 | 0.37 | 0.01 |
| | Max. count: 900 | ✓ | 14 | 1283.6 (39 - 2225) | 0.85 | 1.33 | 0.33 | 0.02 |
| | Max. count: 1000 | ✓ | 15 | 1180.7 (7 - 2431) | 0.85 | 1.32 | 0.35 | 0.01 |
| **GNN4DM** | Max. count: 500 | ✓ | 500 | 120.8 (6 - 659) | 0.99 | 3.86 | 0.72 | -0.00 |
| | Max. count: 600 | ✓ | 600 | 102.3 (4 - 603) | 0.99 | 3.91 | 0.73 | -0.00 |
| | Max. count: 700 | ✓ | 700 | 95.4 (4 - 527) | 0.99 | 4.26 | 0.73 | -0.00 |
| | Max. count: 800 | ✓ | 800 | 81.8 (4 - 666) | 0.99 | 4.17 | 0.75 | -0.01 |
| | Max. count: 900 | ✓ | 900 | 75.0 (2 - 462) | 0.99 | 4.30 | 0.76 | -0.01 |
| | Max. count: 1000 | ✓ | 993 | 72.3 (3 - 544) | 0.99 | 4.57 | 0.76 | -0.01 |
| Pathways | BioCarta | ✓ | 221 | 19.0 (10 - 78) | 0.08 | 3.17 | 0.94 | -0.10 |
| | KEGG | ✓ | 186 | 66.0 (10 - 321) | 0.31 | 2.51 | 0.77 | -0.03 |
| | Reactome | ✓ | 1288 | 53.3 (10 - 459) | 0.63 | 6.93 | 0.83 | -0.01 |
| | WikiPathways | ✓ | 621 | 47.4 (10 - 431) | 0.45 | 4.11 | 0.88 | -0.03 |
| Gene Ontology | Biological process | ✓ | 13915 | 69.7 (1 - 13687) | 0.87 | 70.86 | 0.95 | -0.00 |
| | Molecular function | ✓ | 4526 | 39.6 (1 - 14303) | 0.91 | 12.55 | 0.95 | -0.01 |
| | Cellular component | ✓ | 1834 | 150.5 (1 - 14238) | 0.90 | 19.39 | 0.87 | -0.01 |

Table S3: Descriptive statistics for the investigated module identification methods and pathway databases, filtered to modules with sizes ranging from 2 to 1000.

| | Method | Overlaps | Count of identified modules | Average size (range) | Node coverage | Mean number of modules per gene | Conductance | Modularity overlap |
|---|---|---|---|---|---|---|---|---|
| General topology-based methods | CPM | ✗ | 540 | 27.3 (2 - 830) | 0.93 | 1.00 | 0.50 | 0.07 |
| | Eigenvector | ✗ | 2 | 883.5 (845 - 922) | 0.11 | 1.00 | 0.29 | 0.01 |
| | Greedy Modularity | ✗ | 126 | 10.6 (2 - 244) | 0.08 | 1.00 | 0.35 | 0.28 |
| | Leiden | ✗ | 25 | 225.5 (5 - 956) | 0.36 | 1.00 | 0.17 | 0.19 |
| | Louvain | ✗ | 683 | 17.3 (2 - 834) | 0.75 | 1.00 | 0.47 | 0.13 |
| | Markov Clustering | ✗ | 49 | 111.8 (2 - 868) | 0.35 | 1.00 | 0.43 | 0.11 |
| | Walktrap | ✗ | 566 | 14.5 (2 - 826) | 0.52 | 1.00 | 0.44 | 0.20 |
| | ANGEL | ✓ | 16 | 10.3 (4 - 47) | 0.01 | 1.01 | 0.17 | 0.54 |
| | ASLPAw | ✓ | 369 | 22.4 (2 - 575) | 0.50 | 1.05 | 0.49 | 0.13 |
| | Core Expansion | ✓ | 1974 | 20.5 (2 - 276) | 0.82 | 3.12 | 0.81 | 0.06 |
| | DPClus | ✓ | 2486 | 6.0 (2 - 185) | 0.77 | 1.22 | 0.78 | -0.21 |
| | Graph entropy | ✓ | 3082 | 5.9 (2 - 198) | 0.51 | 2.26 | 0.83 | -0.24 |
| | LFM | ✓ | 289 | 47.7 (2 - 959) | 0.65 | 1.35 | 0.44 | 0.08 |
| | LPANNI | ✓ | 895 | 20.7 (2 - 966) | 0.89 | 1.32 | 0.56 | 0.14 |
| | Node Perception | ✓ | 8693 | 5.7 (2 - 978) | 0.92 | 3.39 | 0.94 | -0.10 |
| | SLPA | ✓ | 227 | 44.3 (2 - 722) | 0.64 | 1.00 | 0.45 | 0.10 |
| DREAM Challenge | DREAM K1 | ✗ | 581 | 27.2 (2 - 100) | 1.00 | 1.00 | 0.59 | 0.08 |
| | DREAM M1 | ✗ | 2029 | 7.8 (1 - 100) | 1.00 | 1.00 | 0.85 | -0.02 |
| | DREAM R1 | ✗ | 327 | 44.7 (3 - 93) | 0.93 | 1.00 | 0.61 | 0.04 |
| UCoDe | Max. count: 500 | ✓ | 137 | 729.5 (5 - 998) | 0.33 | 19.43 | 0.17 | 0.00 |
| | Max. count: 600 | ✓ | 186 | 732.9 (12 - 993) | 0.29 | 30.07 | 0.16 | 0.00 |
| | Max. count: 700 | ✓ | 164 | 793.0 (5 - 1000) | 0.18 | 45.78 | 0.14 | 0.00 |
| | Max. count: 800 | ✓ | 130 | 719.8 (2 - 974) | 0.19 | 31.13 | 0.11 | -0.00 |
| | Max. count: 900 | ✓ | 152 | 725.8 (9 - 993) | 0.21 | 33.09 | 0.11 | 0.00 |
| | Max. count: 1000 | ✓ | 209 | 715.3 (82 - 952) | 0.23 | 40.97 | 0.14 | 0.00 |
| NOCD | Max. count: 500 | ✓ | 5 | 699.2 (437 - 950) | 0.21 | 1.06 | 0.37 | 0.03 |
| | Max. count: 600 | ✓ | 3 | 318.3 (55 - 697) | 0.06 | 1.00 | 0.28 | 0.04 |
| | Max. count: 700 | ✓ | 6 | 769.2 (352 - 980) | 0.26 | 1.11 | 0.34 | 0.03 |
| | Max. count: 800 | ✓ | 5 | 717.0 (13 - 985) | 0.21 | 1.06 | 0.41 | 0.03 |
| | Max. count: 900 | ✓ | 4 | 671.8 (39 - 987) | 0.16 | 1.07 | 0.28 | 0.04 |
| | Max. count: 1000 | ✓ | 6 | 520.0 (7 - 877) | 0.19 | 1.04 | 0.37 | 0.03 |
| **GNN4DM** | Max. count: 500 | ✓ | 500 | 120.8 (6 - 659) | 0.99 | 3.86 | 0.72 | -0.00 |
| | Max. count: 600 | ✓ | 600 | 102.3 (4 - 603) | 0.99 | 3.91 | 0.73 | -0.00 |
| | Max. count: 700 | ✓ | 700 | 95.4 (4 - 527) | 0.99 | 4.26 | 0.73 | -0.00 |
| | Max. count: 800 | ✓ | 800 | 81.8 (4 - 666) | 0.99 | 4.17 | 0.75 | -0.01 |
| | Max. count: 900 | ✓ | 900 | 75.0 (2 - 462) | 0.99 | 4.30 | 0.76 | -0.01 |
| | Max. count: 1000 | ✓ | 993 | 72.3 (3 - 544) | 0.99 | 4.57 | 0.76 | -0.01 |
| Pathways | BioCarta | ✓ | 221 | 19.0 (10 - 78) | 0.08 | 3.17 | 0.94 | -0.10 |
| | KEGG | ✓ | 186 | 66.0 (10 - 321) | 0.31 | 2.51 | 0.77 | -0.03 |
| | Reactome | ✓ | 1288 | 53.3 (10 - 459) | 0.63 | 6.93 | 0.83 | -0.01 |
| | WikiPathways | ✓ | 621 | 47.4 (10 - 431) | 0.45 | 4.11 | 0.88 | -0.03 |
| Gene Ontology | Biological process | ✓ | 10995 | 42.2 (2 - 995) | 0.83 | 35.62 | 0.94 | -0.00 |
| | Molecular function | ✓ | 3081 | 32.1 (2 - 991) | 0.74 | 8.47 | 0.93 | -0.03 |
| | Cellular component | ✓ | 1557 | 48.1 (2 - 945) | 0.64 | 7.44 | 0.87 | -0.06 |

Table S4: Final setting of hyper-parameters in GNN4DM.

| Hyper-parameter | Value |
|---|---|
| Number of GCN layers | 2 |
| Size of hidden GCN layer | 128 |
| Activation functions | (1) ReLU, (2) SoftPlus |
| BCE loss weight ($\lambda$) | 10.0 |
| Optimizer | AdamW |
| Learning rate | 0.001 |
| Learning rate scheduler | Step ($\gamma$: 0.85, step size: 300) |

### 4 Supplementary Figures

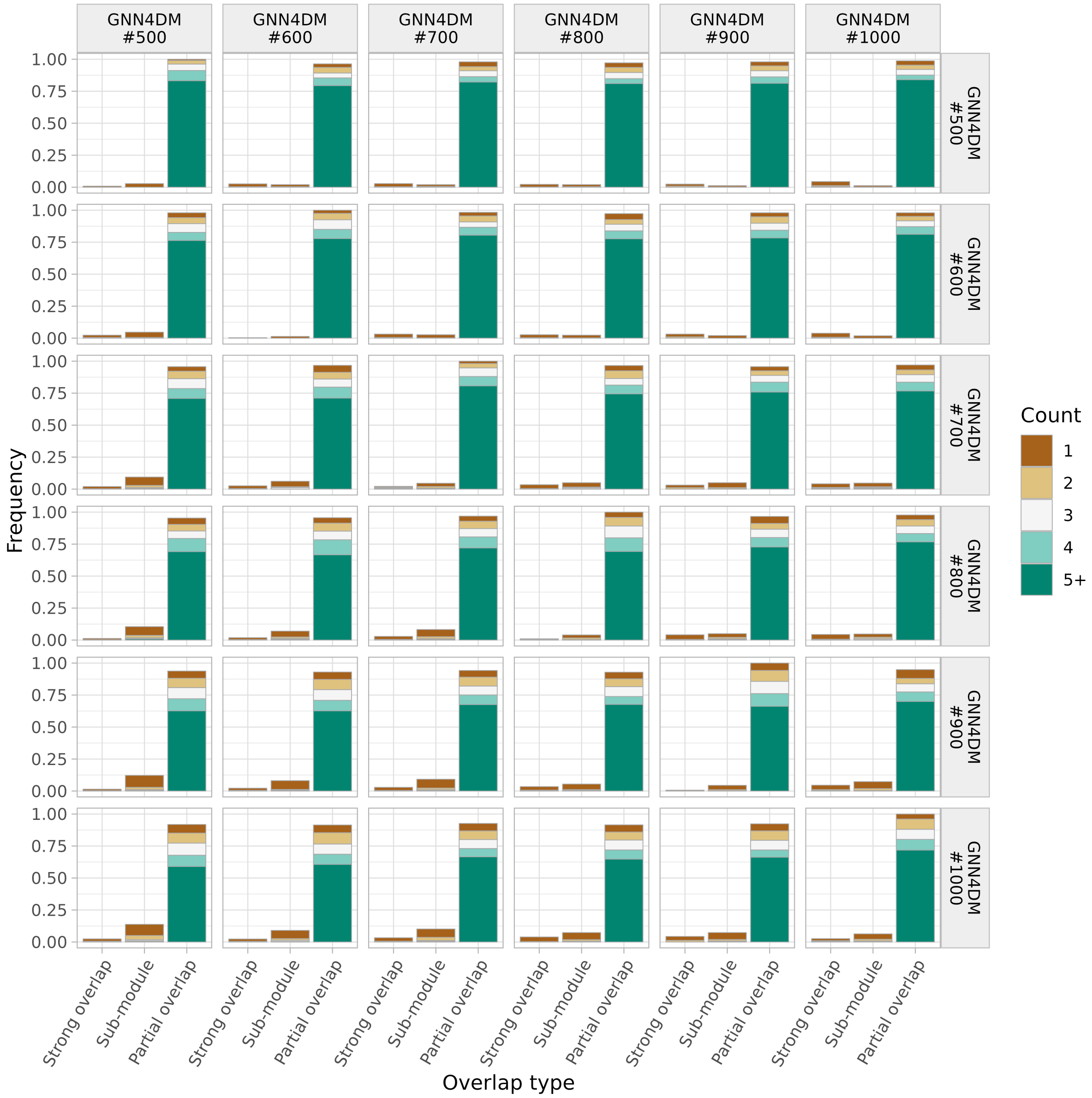


Figure S1: Characterization of all pairwise overlaps between GNN4DM-identified modules. See Supplementary Methods for the definition of the various overlap types.

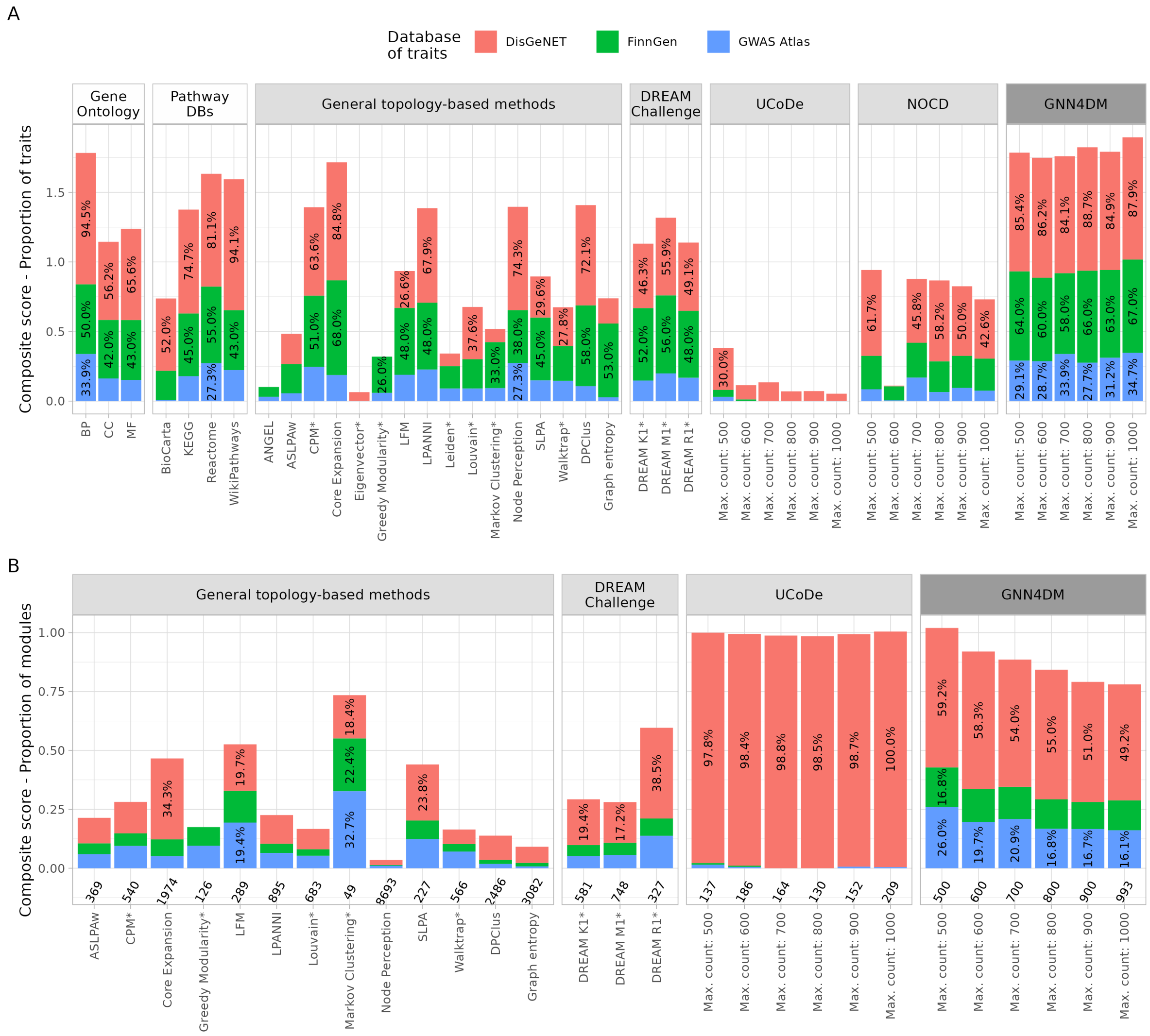


Figure S2: Performance evaluation of module identification methods and pathway databases using significance threshold FDR = 0.01. See the description of Figure 2 for details.

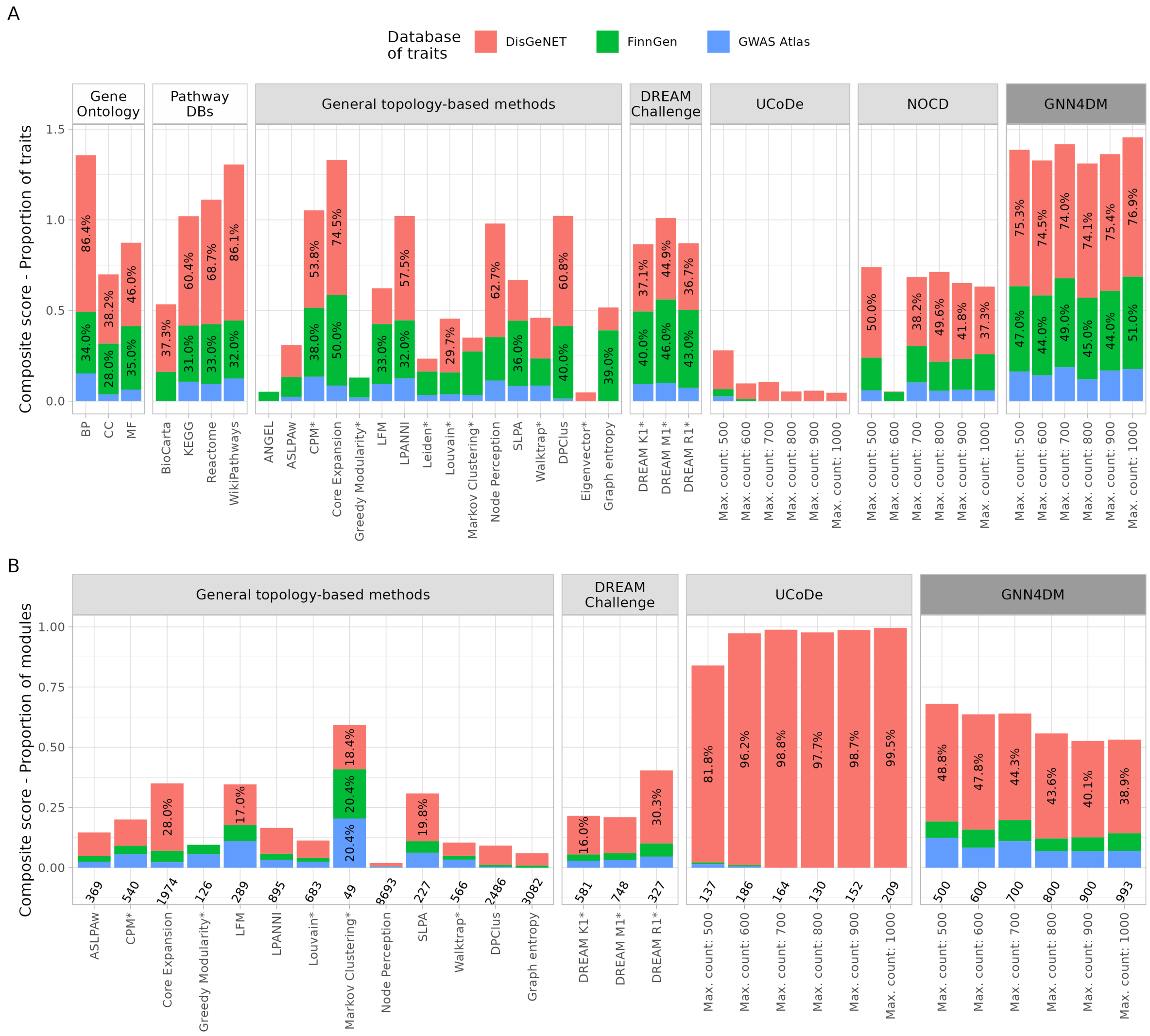


Figure S3: Performance evaluation of module identification methods and pathway databases using significance threshold FDR = 0.001. See the description of Figure 2 for details.

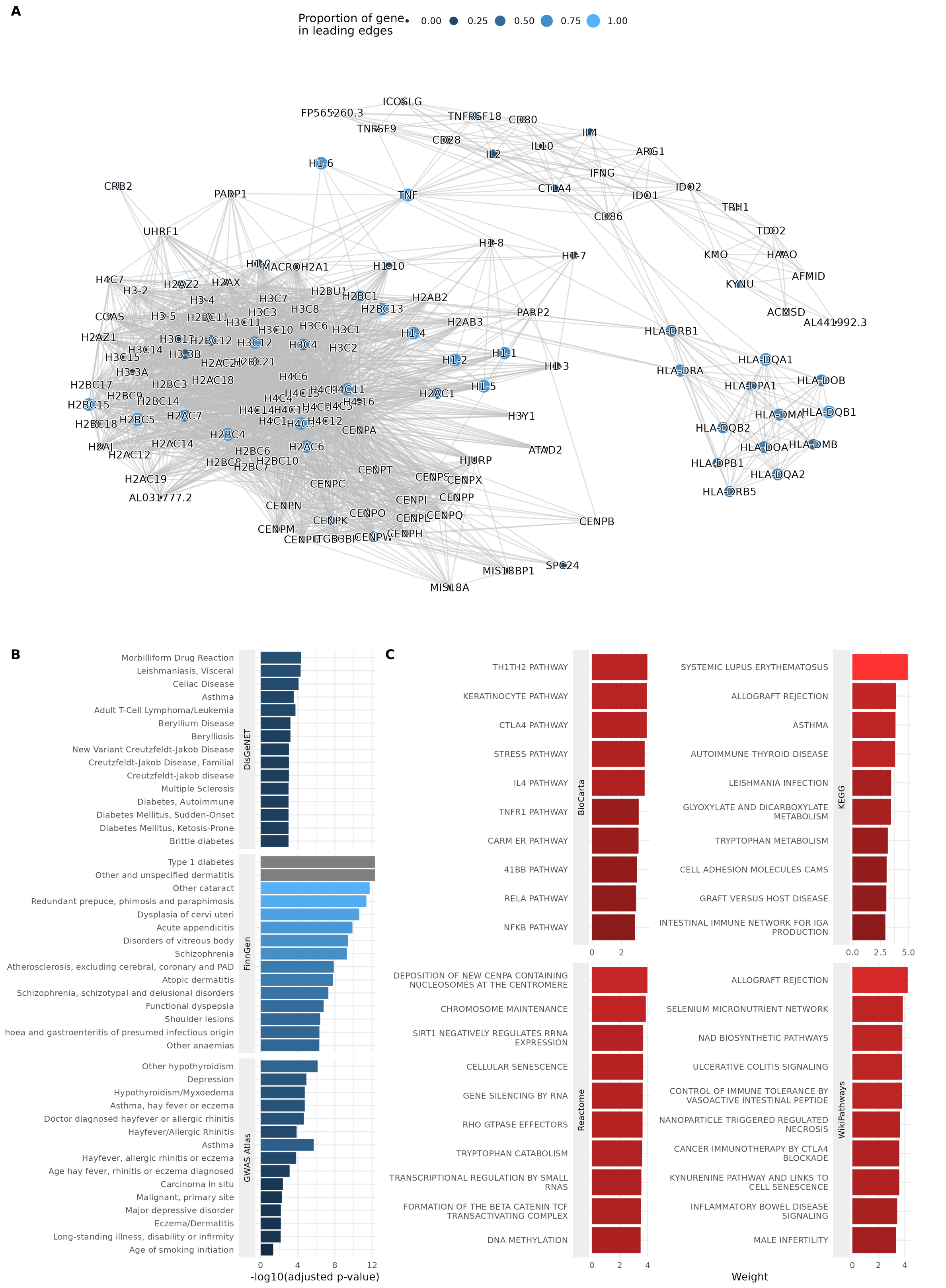


Figure S4: A highly enriched multimorbidity module. (**A**) The graph displays a subnetwork of the STRING PPI network, representing the module. The size and color of each node denote the gene's proportion among the leading edge genes for those diseases in which the module is significantly enriched, as determined by GWAS Atlas. (**B**) The top 15 diseases significantly associated with the module, as identified in the three evaluation datasets. (**C**) The top 10 pathways in each pathway database that are relevant to the module. The bars show the weight GNN4DM learned for the module in association with the corresponding pathway.

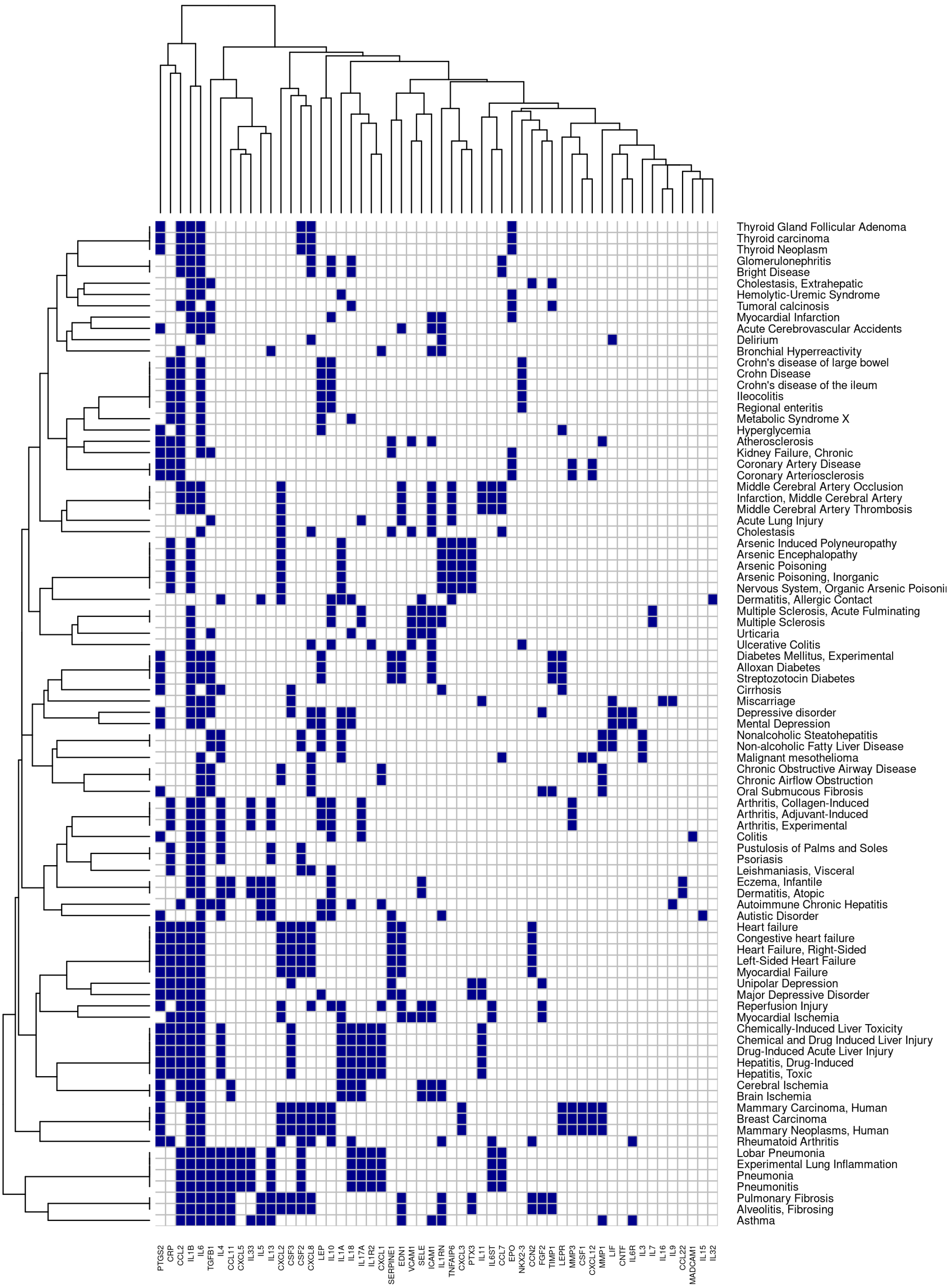


Figure S5: Overlap between genes of the module shown in main text's Figure 3 and the genes related to the disorders significantly associated with the module. Gene-disease associations originate from the DisGeNET database. Only disorders with FDR< 0.0001 and their related genes are shown. Both genes and disorders are hierarchically clustered using Ward's method.

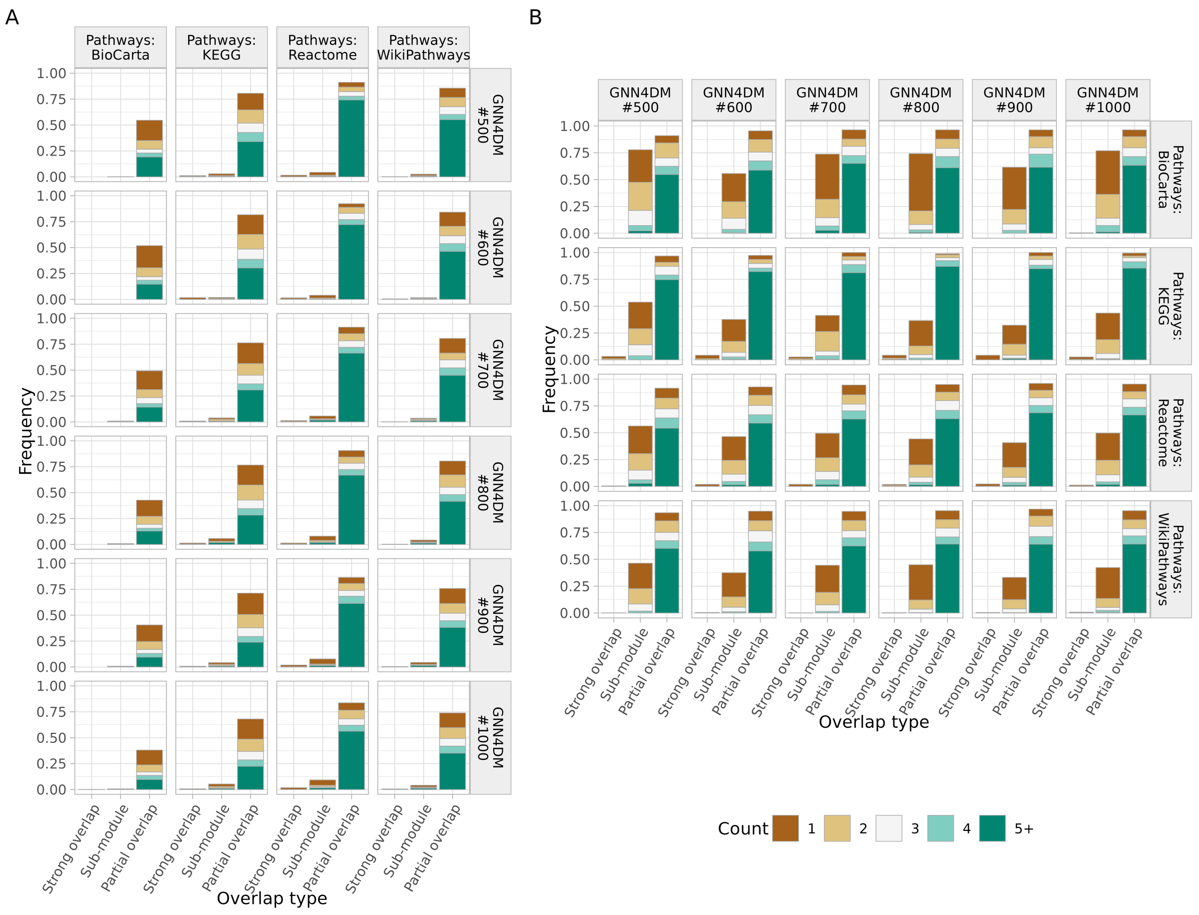


Figure S6: Characterization of all pairwise overlaps between GNN4DM-identified modules and pathway databases. See Supplementary Methods for the definition of the various overlap types.

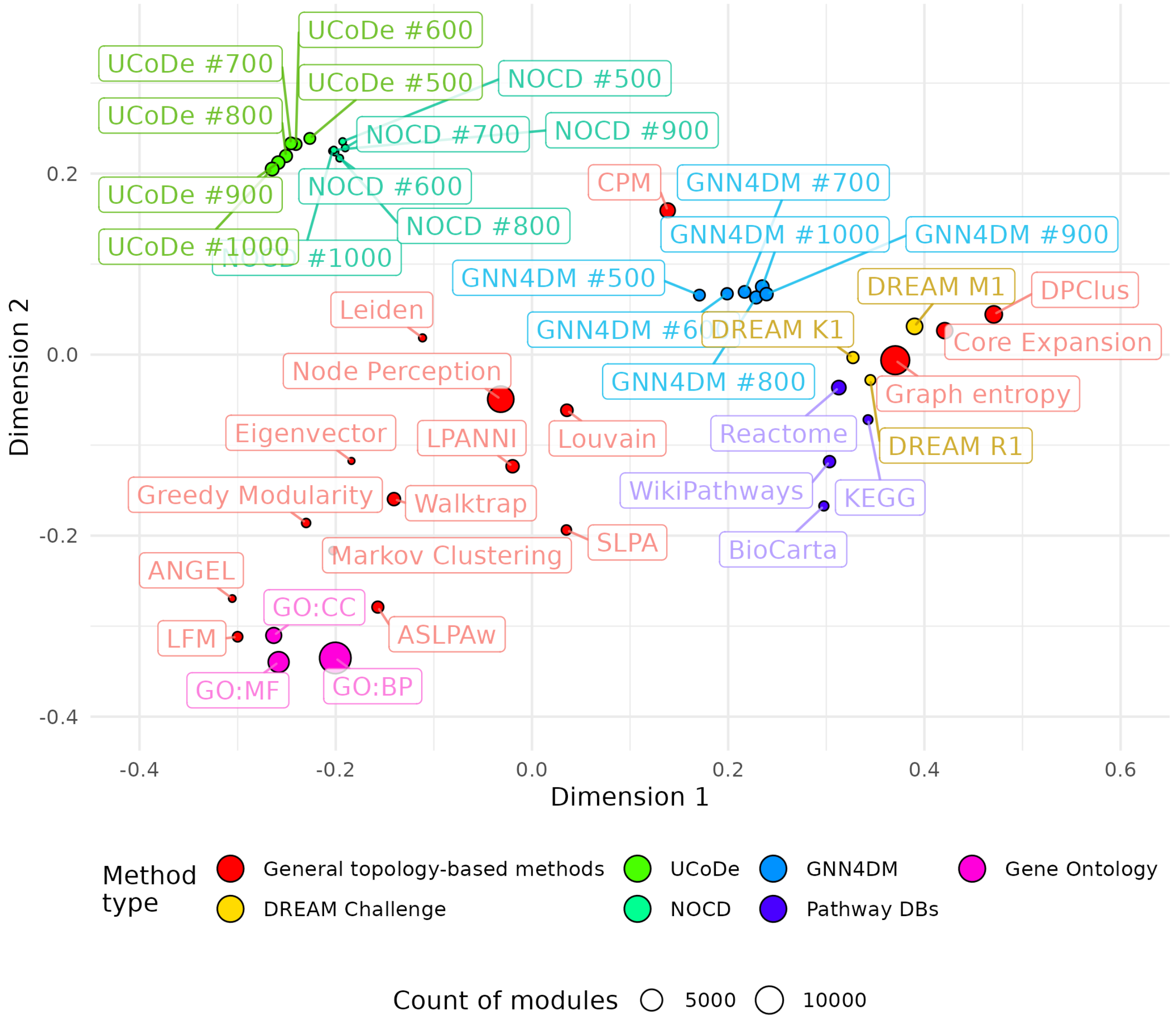


Figure S7: Similarity and complementarity of different module identification methods. Each point corresponds to a module identification method or pathway database. The proximity of two points in the plot indicates a higher similarity in their corresponding module predictions (multidimensional scaling, see Supplementary Methods). The size of the points corresponds to the count of identified modules. The points' and labels' color indicates the type of the methods.

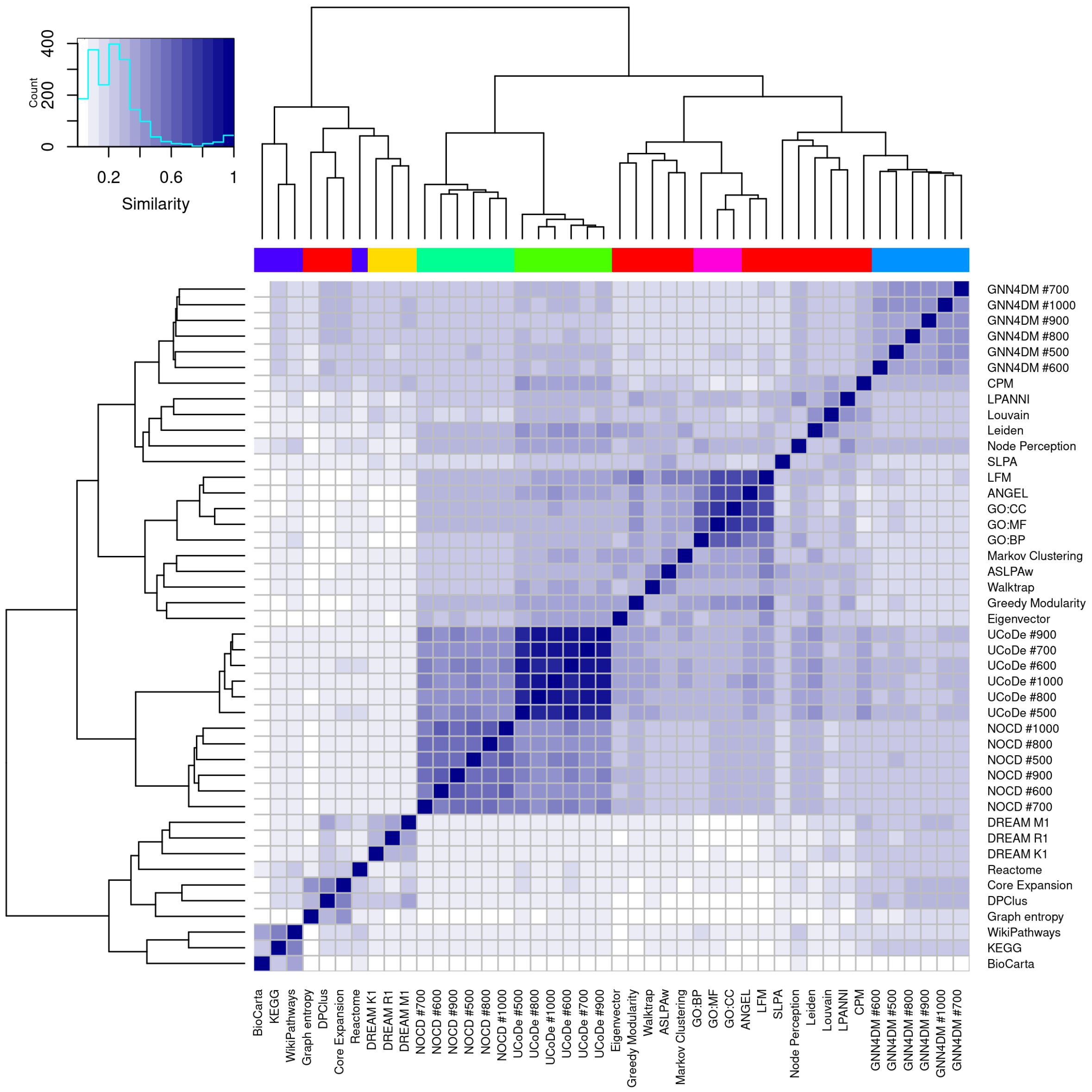


Figure S8: Pairwise similarity of different module identification methods. Similarity was computed based on whether the same genes were clustered together by the two methods. Specifically, a prediction vector was defined for every method, specifying for every gene pair how many modules they were co-clustered in (see Supplementary Methods). A corresponding distance matrix between all methods was computed as the inverse cosine similarity of the prediction vectors and hierarchically clustered using Ward's method. The annotation row indicates the method type.

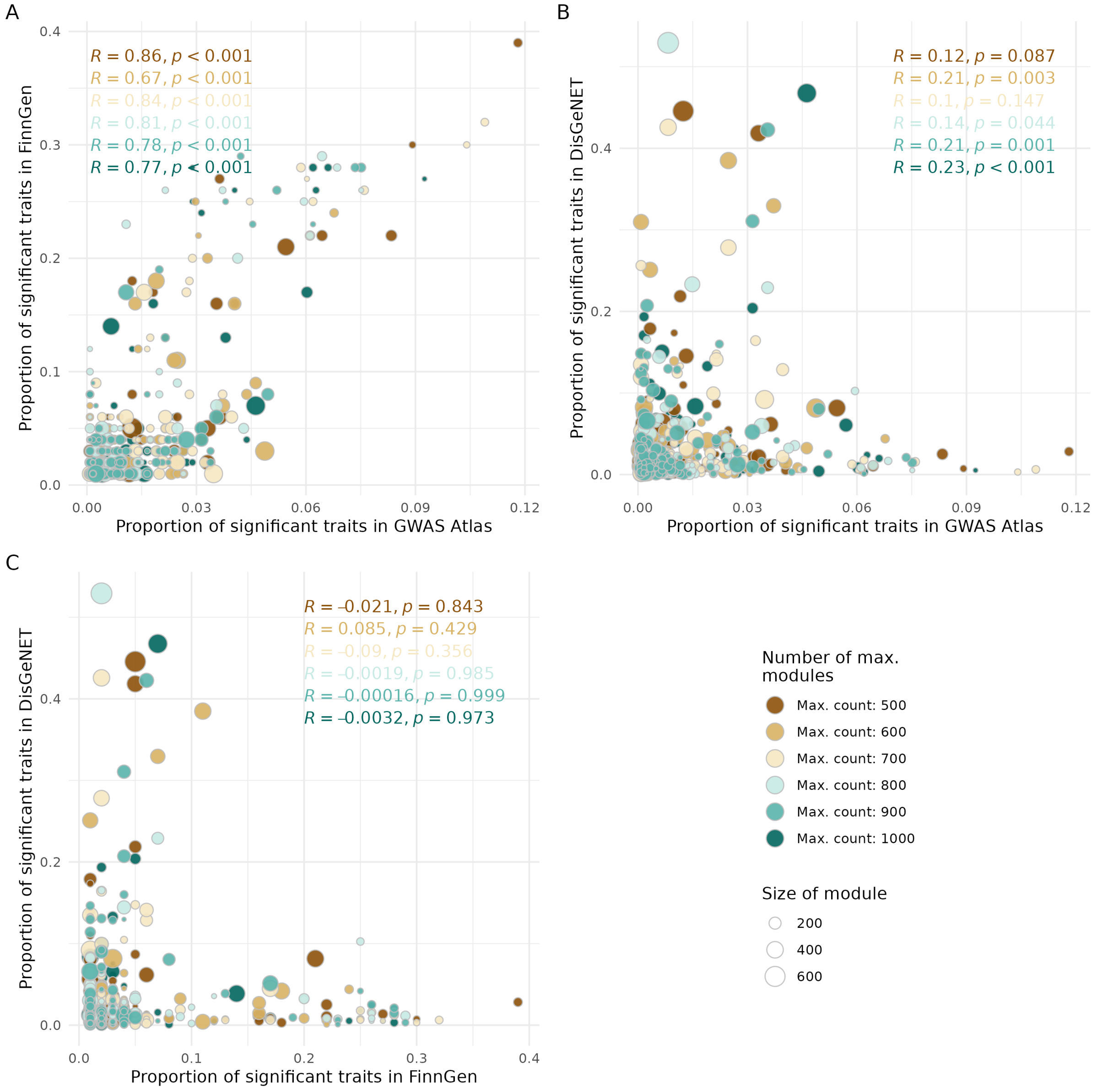


Figure S9: Scatter plots of identified modules according to GNN4DM. Each point corresponds to a module identified by GNN4DM. The color of the nodes denote the maximum module count parameter of the model. The size of the nodes indicates the number of genes assigned to the module. The Pearson correlation coefficient and its associated p-values are shown, indicating the extent of correlation across the evaluation datasets.

### References

[1] Md Altaf-Ul-Amin, Yoko Shinbo, Kenji Mihara, Ken Kurokawa, and Shigehiko Kanaya. Development and implementation of an algorithm for detection of protein complexes in large interaction networks. *BMC Bioinformatics*, 7:207, 12 2006.

[2] Vincent D Blondel, Jean-Loup Guillaume, Renaud Lambiotte, and Etienne Lefebvre. Fast unfolding of communities in large networks. *Journal of Statistical Mechanics: Theory and Experiment*, 2008:P10008, 10 2008.

[3] Sarvenaz Choobdar, Mehmet E. Ahsen, Jake Crawford, Mattia Tomasoni, Tao Fang, David Lamparter, Junyuan Lin, Benjamin Hescott, Xiaozhe Hu, Johnathan Mercer, Ted Natoli, Rajiv Narayan, Aravind Subramanian, Jitao D. Zhang, Gustavo Stolovitzky, Zoltán Kutalik, Kasper Lage, Donna K. Slonim, Julio Saez-Rodriguez, Lenore J. Cowen, Sven Bergmann, and Daniel Marbach. Assessment of network module identification across complex diseases. *Nature Methods*, 16:843–852, 9 2019.

[4] Ali Choumane, Ali Awada, and Ali Harkous. Core expansion: a new community detection algorithm based on neighborhood overlap. *Social Network Analysis and Mining*, 10:30, 12 2020.

[5] Aaron Clauset, M. E. J. Newman, and Cristopher Moore. Finding community structure in very large networks. *Physical Review E*, 70:066111, 12 2004.

[6] A J Enright, S Van Dongen, and C A Ouzounis. An efficient algorithm for large-scale detection of protein families. *Nucleic acids research*, 30:1575–84, 4 2002.

[7] Edward Casey Kenley and Young-Rae Cho. Detecting protein complexes and functional modules from protein interaction networks: A graph entropy approach. *PROTEOMICS*, 11:3835–3844, 10 2011.

[8] Andrea Lancichinetti, Santo Fortunato, and János Kertész. Detecting the overlapping and hierarchical community structure in complex networks. *New Journal of Physics*, 11:033015, 3 2009.

[9] Meilian Lu, Zhenglin Zhang, Zhihe Qu, and Yu Kang. Lpanni: Overlapping community detection using label propagation in large-scale complex networks. *IEEE Transactions on Knowledge and Data Engineering*, 31:1736–1749, 9 2019.

[10] M. E. J. Newman. Finding community structure in networks using the eigenvectors of matrices. *Physical Review E*, 74:036104, 9 2006.

[11] Pascal Pons and Matthieu Latapy. Computing communities in large networks using random walks. In pInar Yolum, Tunga Güngör, Fikret Gürgen, and Can Özturan,

editors, *Computer and Information Sciences - ISCIS 2005*, pages 284–293, Berlin, Heidelberg, 2005. Springer Berlin Heidelberg.

[12] Giulio Rossetti. Angel: efficient, and effective, node-centric community discovery in static and dynamic networks. *Applied Network Science*, 5:26, 12 2020.

[13] Sucheta Soundarajan and John E. Hopcroft. Use of local group information to identify communities in networks. *ACM Transactions on Knowledge Discovery from Data*, 9:1–27, 4 2015.

[14] Mattia Tomasoni, Sergio Gómez, Jake Crawford, Weijia Zhang, Sarvenaz Choobdar, Daniel Marbach, and Sven Bergmann. Monet: a toolbox integrating top-performing methods for network modularization. *Bioinformatics*, 36:3920–3921, 6 2020.

[15] V. A. Traag, P. Van Dooren, and Y. Nesterov. Narrow scope for resolution-limit-free community detection. *Physical Review E*, 84:016114, 7 2011.

[16] Vincent Antonio Traag, Ludo Waltman, and Nees Jan van Eck. From louvain to leiden: guaranteeing well-connected communities. *Scientific Reports*, 9, 2018.

[17] Jierui Xie, Boleslaw K. Szymanski, and Xiaoming Liu. Slpa: Uncovering overlapping communities in social networks via a speaker-listener interaction dynamic process. In *2011 IEEE 11th International Conference on Data Mining Workshops*, pages 344–349, 2011.
